# MitoStructSeg: mitochondrial structural complexity resolution via adaptive learning for cross-sample morphometric profiling

**DOI:** 10.1101/2024.06.28.601295

**Authors:** Xinsheng Wang, Xiaohua Wan, Buqing Cai, Zhuo Jia, Yuanbo Chen, Shuai Guo, Zheng Liu, Fuwei Li, Fa Zhang, Bin Hu

**Affiliations:** Key Laboratory of Brain Health Intelligent Evaluation and Intervention(Beijing Institute of Technology), Ministry of Education, Beijing, China; School of Medical Technology, Beijing Institute of Technology, Beijing, China; Cryo-electron Microscopy Center, Southern University of Science and Technology, Shenzhen, China

**Author notes:** Corresponding authors: Xiaohua Wan, Fa Zhang, Bin Hu.

## Abstract

Mitochondrial morphology and structural changes are closely associated with metabolic dysfunction and disease progression. However, the structural complexity of mitochondria presents a major challenge for accurate segmentation and analysis. Most existing methods focus on delineating entire mitochondria but lack the capability to resolve fine internal features, particularly cristae. In this study, we introduce MitoStructSeg, a deep learning-based framework for mitochondrial structure segmentation and quantitative analysis. The core of MitoStructSeg is AMM-Seg, a novel model that integrates domain adaptation to improve cross-sample generalization, dual-channel feature fusion to enhance structural detail extraction, and continuity learning to preserve spatial coherence. This architecture enables accurate segmentation of both mitochondrial membranes and intricately folded cristae. MitoStructSeg further incorporates a quantitative analysis module that extracts key morphological metrics, including surface area, volume, and cristae density, allowing comprehensive and scalable assessment of mitochondrial morphology. The effectiveness of our approach has been validated on both human myocardial tissue and mouse kidney tissue, demonstrating its robustness in accurately segmenting mitochondria with diverse morphologies. In addition, we provide an open source, user-friendly tool to ensure practical usability.

---

Mitochondria play a central role in cell biology and are considered the powerhouses of the cell^1–3^ due to their critical function in energy production. Understanding the structure of mitochondria is essential to unravel the mechanisms that facilitate efficient energy conversion and regulate biological processes^4–7^. Some research shows that in damaged biological tissue, mitochondrial structure can undergo varying degrees of disruption^8,9^. The structure of mitochondria consists mainly of two membranes: the outer membrane and the inner membrane that is folded into cristae. Furthermore, the cristae damage is strongly associated with numerous diseases^10–12^, highlighting the importance of detailed studies on mitochondrial structure. Given mitochondria complex subcellular structure, detailed imaging techniques are required to fully understand mitochondria morphology and function. Scanning electron microscopy (SEM)^13,14^ is commonly employed for this purpose, offering high-resolution images of mitochondrial ultrastructure. In particular, focused ion beam scanning electron microscopy (FIB-SEM)^15–18^ offers several distinct advantages, allowing for three-dimensional reconstruction with isotropic resolution in the low nanometer range. These capabilities make FIB-SEM particularly effective for examining the fine details of mitochondrial structure. However, the complex and variable morphology of mitochondria^19,20^ presents significant challenges for image segmentation^21–26^. The image noise and the variability in mitochondrial size and shape complicate segmentation. Additionally, the intricate folding of the inner membrane and its proximity to the outer membrane, along with the varying shapes and densities of cristae, make accurate segmentation particularly challenging.

The widespread adoption of deep learning technologies across various fields has highlighted its growing potential in biomedical image analysis^27–29^. DeepContact^30^ is a UNet-based^31^ model designed to segment and analyze membrane contact sites between mitochondria and the ER in electron microscopy images. However, it cannot segment the mitochondrial cristae, preventing the accurate capture of the detailed mitochondrial structure and morphological changes. PHILOW^32^ utilizes a HITL-TAP model to automatically segment the topological structure of mitochondrial cristae. However, it requires manual interaction to refine and correct the model’s output, limiting its fully automated capability. Moreover, most existing methods primarily focus on generating initial segmentation results and lack quantitative analysis of these results^30,33–35^. Quantitative analysis of mitochondrial information is crucial for understanding their dynamics and alterations in disease contexts, as it provides insights into mitochondrial morphology and health status. To better understand mitochondrial changes during disease progression, there is an urgent need for a tool that can not only accurately segment mitochondrial structure but also provide quantitative analysis of the segmentation outcomes.

To address this problem comprehensively, we have developed MitoStructSeg, the first fully automated framework that both integrates mitochondrial structure segmentation and quantitative analysis. Our framework includes the Adaptive Multi-domain Mitochondrial segmentation module (AMM-Seg), a novel algorithm we designed that automatically segments both the mitochondrial outer membrane and inner cristae. In addition, MitoStructSeg provides advanced tools for quantifying key mitochondrial features such as volume, surface area, and cristae density. Analyzing these results, our method explores correlations between mitochondrial characteristics and health status, providing valuable insights into mitochondrial function and pathology. To evaluate our method’s performance on mitochondria with structural diversity, we validate MitoStructSeg on samples from human myocardial tissue and mouse kidney tissue, demonstrating its effectiveness in handling diverse mitochondrial morphologies.

## Results

### The design of MitoStructSeg

MitoStructSeg is an integrated framework that automatically segments mitochondrial structures, including the outer membrane and inner cristae, and quantitatively analyzes the segmentation results to identify mitochondrial damage. As shown in the Fig.1, the MitoStructSeg consists of two modules: the mitochondrial structure segmentation module and the mitochondrial quantitative analysis module. The mitochondrial structure segmentation module is based on the Adaptive Multidomain Mitochondrial segmentation (AMM-Seg) network architecture (Supplementary Fig.1) that we proposed. The mitochondrial quantitative analysis module processes segmentation results to generate quantitative data, including surface area, volume, and cristae density. In addition, we perform 3D visualization^36^ of the segmentation results to further illustrate the structural details and highlight the state of mitochondrial health and damage.

**Fig. 1.**
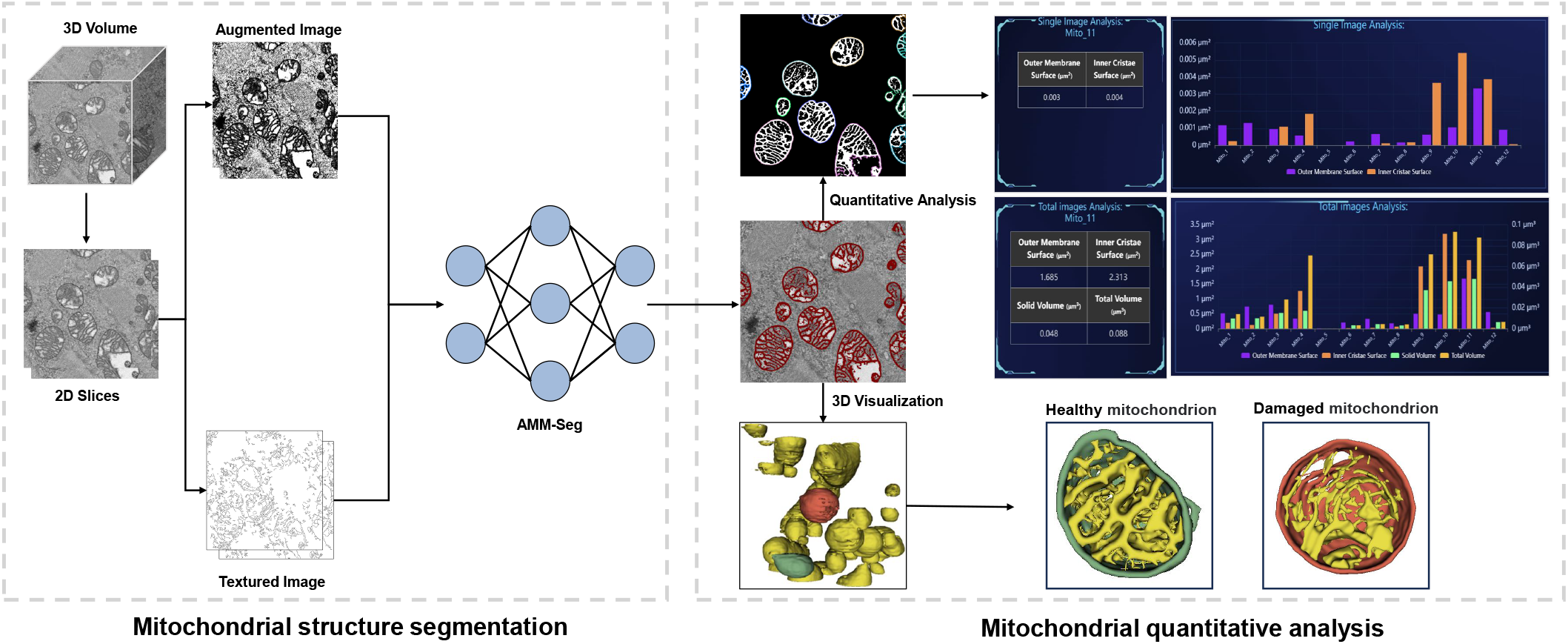
The architecture of MitoStructSeg. MitoStructSeg’s workflow includes mitochondrial structure segmentation and mitochondrial quantitative analysis. **Mitochondrial structure segmentation,** First, 3D mitochondrial volume blocks are sliced to generate 2D slices. The 2D slices are then processed to generate augmented images, which enhance feature diversity, and texture images, which provide detailed texture features. The augmented images and texture images are subsequently input into the AMM-Seg, yielding segmentation results of the mitochondrial structure. **Mitochondrial quantitative analysis,** A user-friendly GUI is provided for statistical analysis of the segmentation results, enabling the extraction of key metrics such as surface area, volume, and cristae density. Additionally, we leverage the open-source tool 3D Slicer for visualizing the segmentation results, clearly illustrating the structural differences between healthy and damaged mitochondria.

The Human Myocardium dataset is obtained from the myocardial tissues of three COVID-19 patients and is collected using FIB-SEM, with a voxel size of 5 nm*×*5 nm and a slice interval of 10 nm (see Methods Section 1 for details). The AMM-Seg model uses 200 labeled images (800 × 800 pixels) from Patient#1 dataset as the source domain, while 200 unlabeled images (800 × 800 pixels) are employed as the target domain for training. The target domain is derived from other datasets to more effectively evaluate the model’s generalization ability on different types of dataset. We focus on the data within the red dotted box (800 × 800 × 800 voxels) for predictive analysis. Additionally, other methods are also based on the same dataset to evaluate results and make comparisons. The segmentation results of AMM-Seg and Ground Truth are visualized to evaluate the actual effect of segmentation (Fig.2a). We utilize three distinct metrics for evaluation: (1) The degree to which the mitochondrial structure in a predicted domain overlaps with a true domain is measured via the Intersectover-Union (IoU) between the mitochondrial structure in the predicted and ground-truth domains. (2) The Matthews Correlation Coefficient (MCC) takes into account all four classification outcomes: true positives, false positives, true negatives, and false negatives. The score ranges from −1 to 1, where 1 signifies perfect prediction, 0 indicates random effects, and −1 denotes complete errors. Since the MCC is unaffected by the uneven distribution of samples, it is particularly well-suited for datasets with class imbalances. (3) The F1 score, a harmonic mean of precision and recall, combines precision and recall to assess the model’s accuracy in identifying true positives.

**Fig. 2.**
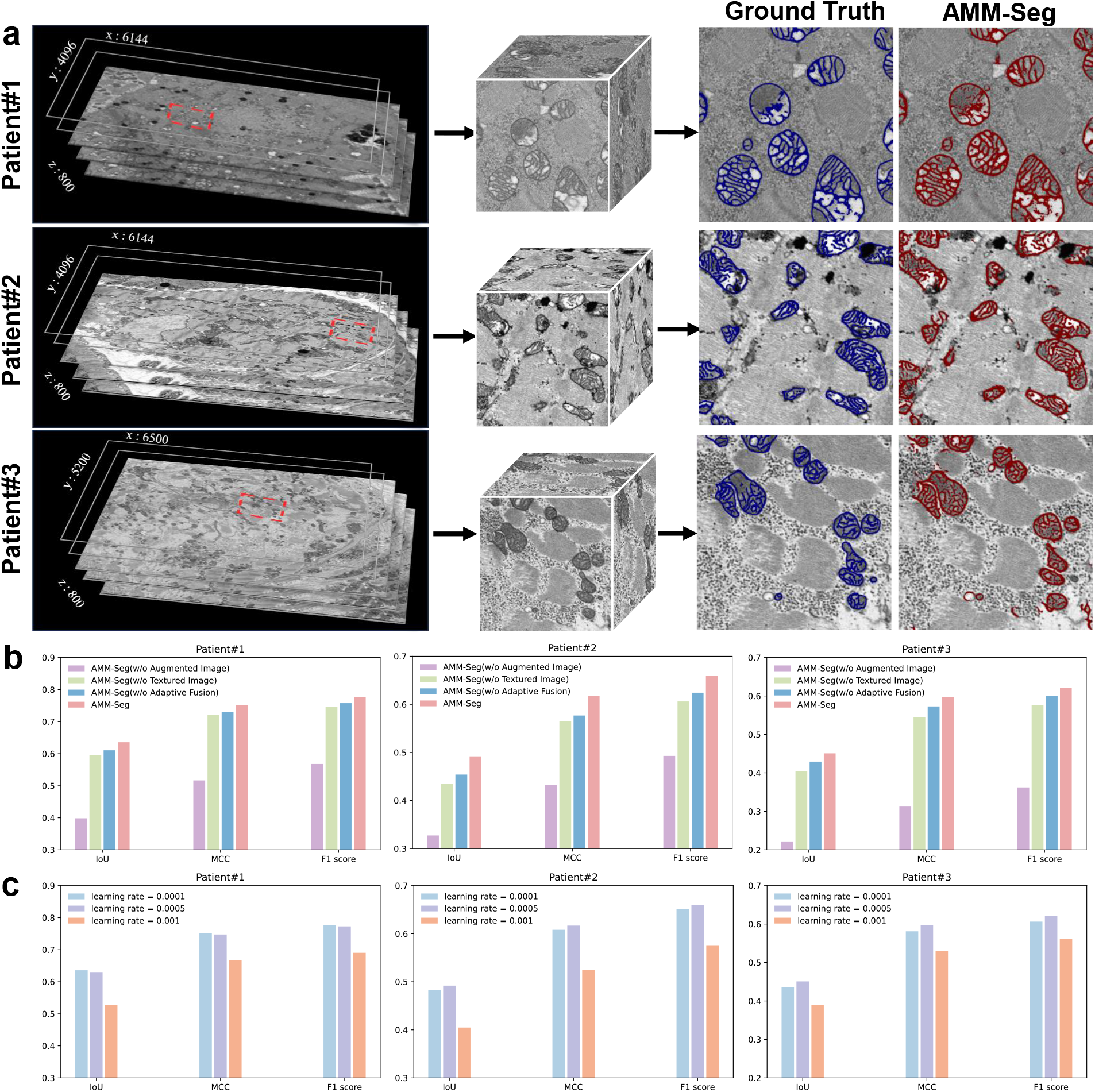
The segmentation performance of AMM-Seg in the Human Myocardium dataset. **a,** The image processing and segmentation results of three human myocardium patient datasets are shown. The left image of each dataset shows the FIB-SEM acquired data, the middle image shows the cropped area indicated by the red dashed box, and the right image compares the ground truth (blue contours) with the segmentation result of the AMM-Seg model (red contours). **b,** The results of the ablation experiments for AMM-Seg are presented. The ablation experiment compares the performance of the AMM-Seg without “Augmented Image”, “Textured Image”, “Adaptive Fusion”, and with the complete AMM-Seg methods. **c,** We compared the effects of different learning rates (0.0001, 0.0005, and 0.001) on the segmentation performance of three human myocardium patient datasets. The results show that different learning rates have a significant impact on segmentation performance. For Patient#1 dataset, a learning rate of 0.0001 yields the best effect, whereas for Patient#2 and Patient#3 datasets, a learning rate of 0.0005 proves most effective.

We perform an ablation study to analyze the role of each component in the proposed model AMM-Seg. In particular, we have the following variants: (1) w/o augmented image: removing the augmented image and only used the textured image. (2) w/o textured image: similar to w/o augmented Image, only the augmented image are used. (3) w/o adaptive fusion: the image features generated by the augmented image and the textured image are fused in equal weight fusion to replace the adaptive fusion module (see Methods Section 2 for details). It can be observed from the Fig.2b that when either the augmented or the textured image is removed, the effect on the three human myocardium patient datasets decreases. This highlights that the model achieves better performance when both the textured image and the augmented image are applied together, outperforming the use of either one alone. This demonstrates that incorporating texture information can effectively enhance the structural features of the image. Finally, an adaptive fusion strategy is employed to integrate the augmented image and the textured image, yielding results significantly superior to those of the equal weight fusion strategy. This indicates that the integration of adaptive fusion mechanism allows the model to dynamically adjust its focus based on the feature’s importance, particularly enhancing its attention towards the structural features of mitochondria. In general, each component contributes to the performance of the proposed model, and their combination shows the best performance.

After validating the effectiveness of the AMM-Seg architecture, we further investigated the impact of optimization strategies on model performance, with a particular focus on learning rate configuration and dynamic scheduling. In deep learning, the learning rate plays a crucial role in determining both the convergence speed and the generalization ability of the model. To ensure robust and stable training, AMM-Seg employs a multi-stage dynamic learning rate schedule, where a specified initial learning rate is progressively reduced using a polynomial decay strategy. This approach allows the model to maintain strong learning capacity during the early training stages while gradually lowering the update rate as training progresses, thereby reducing the risk of overfitting and supporting stable convergence. As illustrated in Fig.2c, we evaluated the model’s performance under this dynamic schedule using three different initial learning rates. The results show that the optimal learning rate varies across datasets: 0.0001 yielded the best performance on the Patient#1 dataset, while 0.0005 performed better for Patient#2 and Patient#3. These findings suggest that the optimal initialization of the learning rate is influenced by the specific structural and textural characteristics of each dataset and, ideally, should be selected adaptively. However, in practice, when prior knowledge about the dataset is limited, an initial learning rate of 0.0005 serves as a robust and effective default across diverse data sources. More importantly, our experiments demonstrate that a well-designed dynamic learning rate strategy plays a critical role in enhancing both the accuracy and stability of the segmentation model, highlighting the importance of coupling architectural design with training optimization for improved performance.

### Comparison of mitochondrial structure segmentation with state-of-the-art methods

We conduct a comparative analysis of the segmentation performance of AMM-Seg against five benchmark methods, including MEDIAR^33,34^, DA-ISC^35^, UNet^31^, Swin unetr^37^ and Unetr^38^. These benchmark methods utilize advanced segmentation techniques, incorporating multimodal data processing, domain adaptation strategies, and novel neural network architectures to improve accuracy and performance in biomedical image analysis. The segmentation results of each method are visually compared, as illustrated in Fig.3a. The UNet, Swin unetr, and Unetr methods perform the poorest among the methods evaluated. Their segmentation results often fail to capture the complete structure of mitochondria, resulting in sparse and blurred contours with high false-positive rates. This demonstrates their inability to effectively segment mitochondrial structure, particularly in datasets with highly complex features. The DA-ISC and MEDIAR methods demonstrate noticeable improvements over UNet, Swin unetr, and Unetr. Their segmentation results are closer to the Ground Truth and manage to capture most structural contours accurately. However, in regions with intricate inner cristae structures, their performances begin to show limitations, with slight deviations from the true contours. In contrast, AMM-Seg demonstrates superior performance in preserving intricate inner cristae structures, consistently achieving segmentation results that closely align with the Ground Truth. Its red contours closely match the blue contours of the Ground Truth, even in the most structurally complex regions. This demonstrates AMM-Seg’s exceptional capability to accurately segment mitochondrial structure and adapt to the complexity of different datasets. Subsequently, to evaluate the effects of all methods more accurately, F1 scores are assessed for the 40 images (800 × 800 pixels) in the validation dataset (Fig.3b). The UNet, Swin unetr, and Unetr methods exhibit the lowest F1 scores, with values that are highly scattered and show significant fluctuations. This reflects their poor stability and limited ability to handle the variability in mitochondrial structure across different datasets. Although DA-ISC and MEDIAR achieve higher median F1 scores than UNet, Swin unetr, and Unetr, their results still show fluctuations across different datasets, indicating that there is still a need for improvement in domain adaptation. AMM-Seg outperforms all other methods, achieving the highest median F1 score with minimal fluctuation across the three human myocardium patient datasets. This indicates not only its superior segmentation accuracy but also its remarkable stability. The method consistently performs well, regardless of the complexity or variability present in the datasets. It is worth noting that most mitochondrial segmentation tasks currently use block segmentation, which has relatively high overall performance metrics^33–35^. For comparison, we also conduct experiments on the mitochondrial block segmentation task and present the results in Fig.4. Compared to mitochondrial block segmentation, mitochondrial structure segmentation is considerably more challenging, as the structures exhibit fine, line-like features that are highly sensitive to minor segmentation errors. Consequently, mitochondrial structure segmentation task yields relatively lower overall performance metrics.

**Fig. 3.**
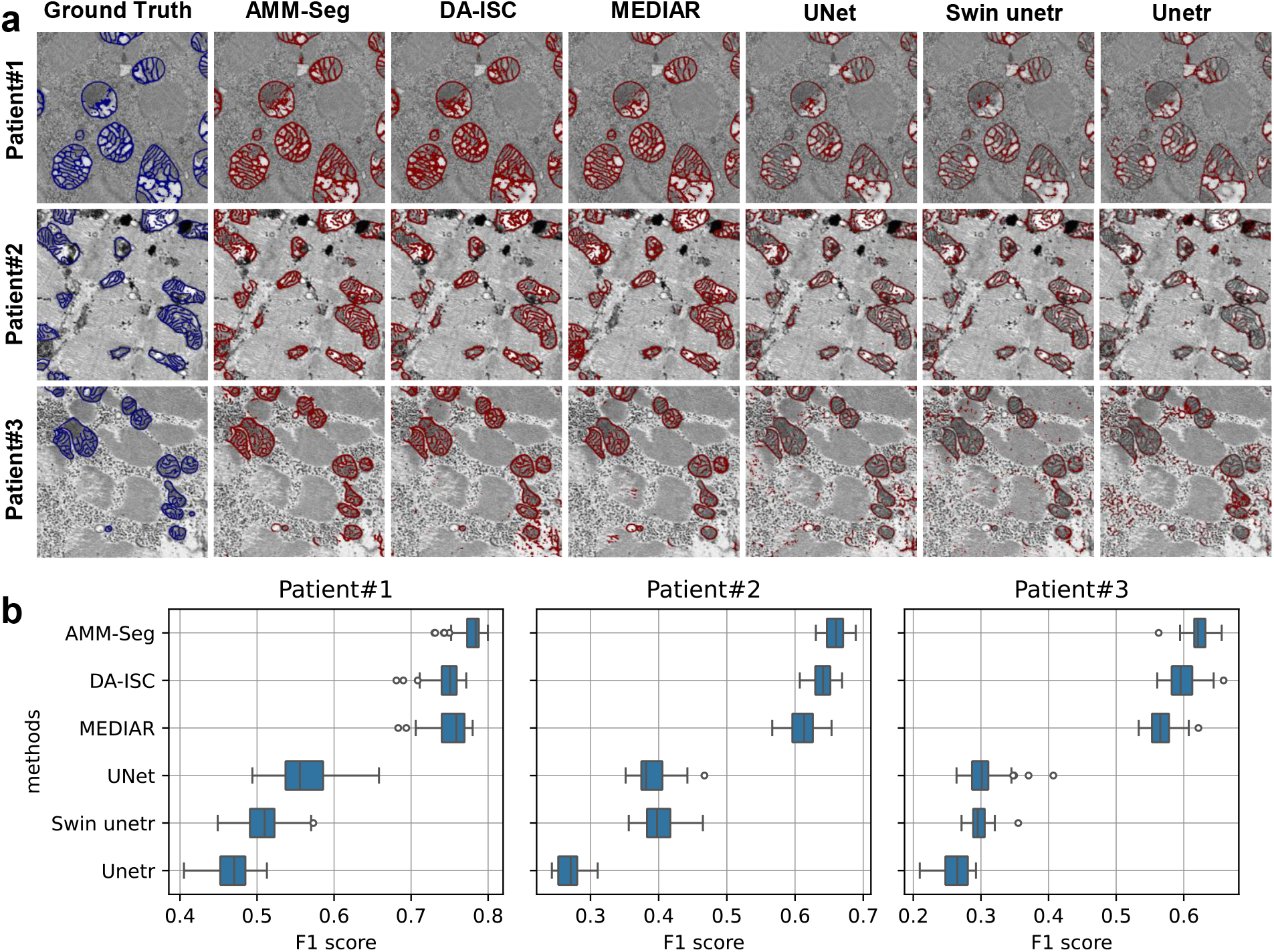
Comparison of AMM-Seg and mainstream methods for structure segmentation in the Human Myocardium dataset. **a,** UNet, Swin unetr and Unetr perform poorly in the three human myocardium patient datasets, producing sparse segmentation results and high false positive rates. The segmentation results of DA-ISC and MEDIAR are relatively close to the true values, but still have some bias when dealing with complex inner cristae structure. In contrast, AMM-Seg method shows superior performance, with its red contours being highly consistent with the ground truth (blue contours), demonstrating an excellent ability to adapt to complex structure. **b,** Quantitative comparison results reveal that the F1 score distributions of the UNet, Swin unetr, and Unetr methods are relatively scattered, showing large fluctuations and lower medians. The DA-ISC and MEDIAR methods perform better, with scores second only to AMM-Seg. Notably, the AMM-Seg method achieves the highest median and the smallest score fluctuation in F1 scores across the three human myocardium patient datasets, demonstrating not only accurate but also highly stable segmentation performance.

To verify the model’s performance on images with different resolutions, we conduct a downsampling experiment using the Human Myocardium dataset. We select representative original images and progressively reduce their resolution to 1/2 and 1/4 of the original size to simulate varying resolution levels. Segmentation predictions are then performed and analyzed at each resolution to evaluate the model’s robustness under different imaging conditions. During the image reduction process, we choose the LANCZOS filter^39^, which effectively reduces the aliasing effects that may be introduced by the reduction operation. Subsequently, to ensure processing consistency, we use nearest-neighbor interpolation to restore all reduced images to a unified 800 × 800 pixels resolution. We then use our previously trained model to make segmentation predictions and evaluate its performance across images of varying resolutions. From the results (Supplementary Fig.2), it is observable that although the image details gradually blur due to the reduction in resolution, the prediction output of the model does not show significant degradation. The AMM-Seg continues to accurately identify and segment the outer membrane and inner cristae of mitochondria. This consistency demonstrates the high generalization ability of our model in processing images of different resolutions.

To further assess the effectiveness and adaptability of AMM-Seg beyond pathological human myocardial tissue, we evaluate its performance on a distinct dataset comprising healthy mouse kidney tissue. We use three evaluation metrics (IoU, MCC, and F1 score) to provide a comprehensive assessment of segmentation performance. As shown in Supplementary Fig.3, AMM-Seg achieves higher visual consistency with the ground truth and outperforms other methods across all three metrics, while maintaining low variability. These results further support the model’s ability to accurately segment mitochondria with diverse structural characteristics.

### Comparison of mitochondrial outer membrane and inner cristae segmentation with state-of-the-art methods

The mitochondrial outer membrane is a closed boundary, while the inner cristae are densely packed and exhibit complex morphology. To evaluate the model’s segmentation performance on these distinct substructures, we conduct separate experiments for outer membrane and inner cristae segmentation, respectively.

The outer membrane segmentation task emphasizes the accurate delineation of smooth, continuous membrane boundaries, which is crucial for evaluating the model’s ability to capture fine structural contours. To assess performance on this task, we conduct a comparative analysis from two perspectives: the traditional mitochondrial block segmentation task and a dedicated outer membrane segmentation task. For the AMM-Seg, DA-ISC, and MEDIAR methods, there is no need to retrain the model specifically for mitochondrial block segmentation. Based on the mitochondrial structure segmentation results from the previous step, the segmentation of mitochondrial blocks can be obtained using the Flood Fill method^40^. However, for the UNet, Swin unetr, and Unetr methods, the segmented outer membranes are often fragmented and structurally discontinuous, making the Flood Fill method inapplicable. Consequently, it is necessary to retrain these models specifically for the mitochondrial block segmentation task. The visualization of the mitochondrial block segmentation results is illustrated in Fig.4a, where green represents the correctly segmented areas, red indicates the missed segments, and blue highlights the erroneously segmented areas. The results show that on Patient#1 dataset, most methods successfully segment the mitochondrial blocks. Moreover, on Patient#2 and Patient#3 datasets, AMM-Seg outperforms other methods in terms of missed and incorrect segmentations. In addition, the F1 score is utilized to evaluate the 40 images in the validation dataset. According to the box plot results in Fig.4b, AMM-Seg achieves the highest performance on the three human myocardium patient datasets, especially on Patient#2 and Patient#3 datasets. This demonstrates that AMM-Seg possesses strong cross-domain capabilities, maintaining effective segmentation across diverse datasets. We also perform mitochondrial block segmentation on the Mouse Kidney dataset. As shown in Supplementary Fig.4, AMM-Seg produces more accurate and consistent results than other methods, with fewer missed and incorrect regions. In addition to the F1 score used previously, we also report IoU and MCC to provide a more comprehensive evaluation. AMM-Seg achieves the highest scores across all three metrics with low variability, demonstrating its effectiveness in handling structurally diverse mitochondria.

**Fig. 4.**
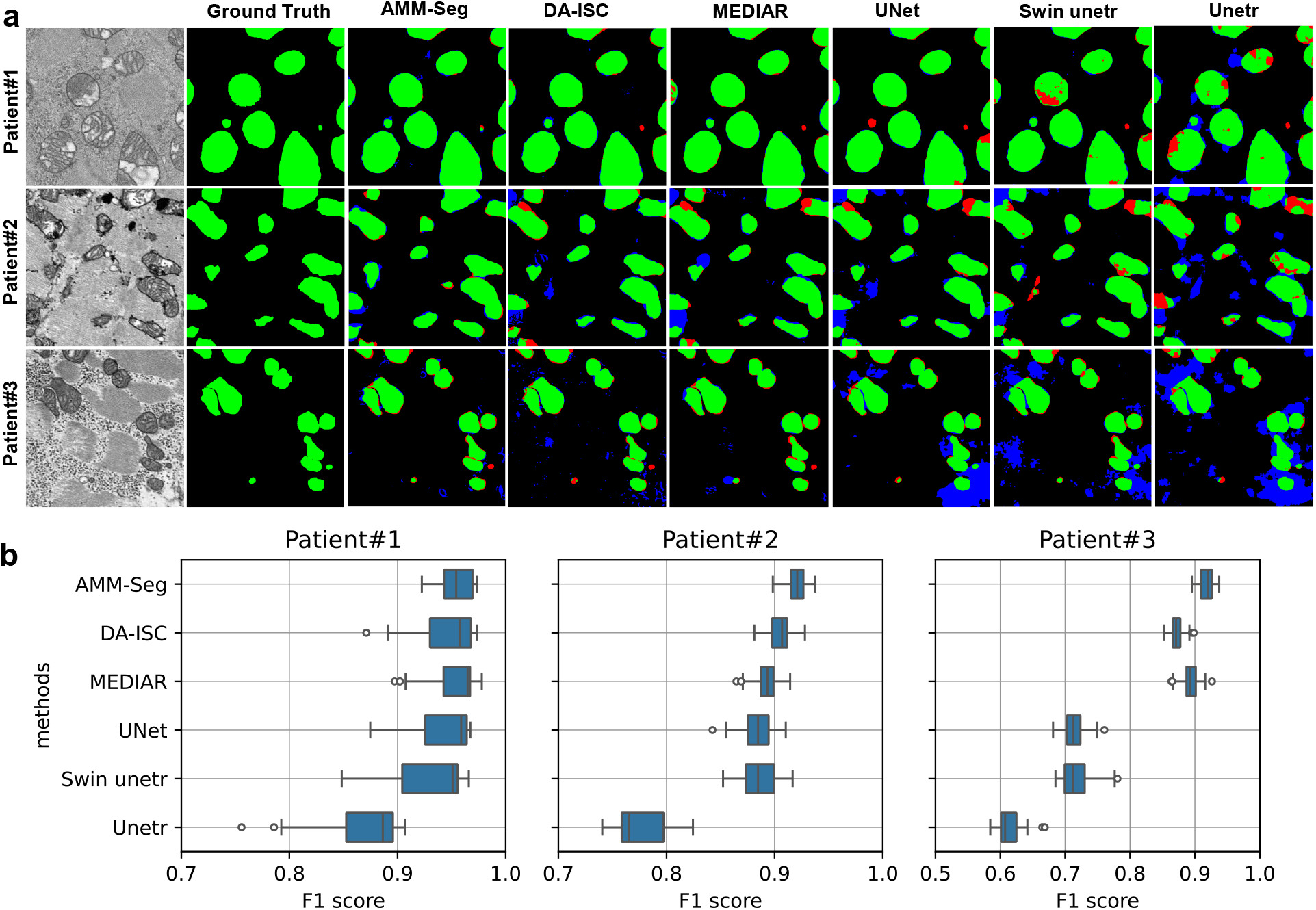
Comparison of AMM-Seg and mainstream methods for block segmentation in the Human Myocardium dataset. **a,** The results of AMM-Seg, DA-ISC, and MEDIAR are obtained by Flood Fill based on the mitochondrial structure segmentation results, while UNet, Swin unetr, and unetr are retrained models for block segmentation tasks. From the visualization results, we can observe that in the Patient#1 dataset, the correct part predicted by AMM-Seg (green block) is significantly more consistent with the Ground Truth, and the missing predicted part (red block) and the wrong predicted part (blue block) are significantly fewer than with other methods. In the Patient#2 and Patient#3 datasets, due to the domain adaptation strategy adopted by AMM-Seg, the erroneous segmentation results (blue block) are significantly less than other methods. **b,** In this experiment, we compare the performance of the AMM-Seg F1 score with that of five other methods on three human myocardium patient datasets. From the results of the box plot, the overall performance of AMM-Seg, DA-ISC, MEDIAR, and UNet in the Patient#1 dataset is similar, but the performance of AMM-Seg is more stable. In the Patient#2 and Patient#3 datasets, AMM-Seg’s performance is significantly better than that of other methods and demonstrates higher stability. This further illustrates the advantage of AMM-Seg in its domain adaptation capabilities.

Furthermore, we retrain all models and conduct segmentation experiments specifically on the mitochondrial outer membrane. On the Human Myocardium dataset (Supplementary Fig.5) and the Mouse Kidney dataset (Supplementary Fig.6), AMM-Seg consistently achieves the best performance. It is worth noting that this experiment focuses on the segmentation of the mitochondrial outer membrane, which exhibits line-like structures, resulting in relatively lower overall evaluation metrics. Compared to the previous block segmentation results, the overall evaluation metrics for outer membrane segmentation are lower. This difference in evaluation metrics can be attributed to the nature of the segmentation targets. Block segmentation typically involves larger, enclosed mitochondrial regions with relatively stable spatial continuity, where small segmentation inaccuracies have limited impact on overall performance scores. In contrast, outer membrane segmentation targets thin, continuous boundary structures that are highly sensitive to edge integrity, where even slight discontinuities or omissions can cause a significant drop in evaluation metrics.

In the inner cristae segmentation task, the focus is on capturing the complex and irregular morphology of the inner cristae, which presents significant challenges due to their intricate structures and high variability. To evaluate the performance of various segmentation approaches, we restrict the labeling to the inner cristae region and retrain all models for a comparative analysis. To verify the performance of AMM-Seg in the task of inner cristae segmentation, we conduct comparative experiments with a variety of mainstream methods. On the Human Myocardium dataset (Supplementary Fig.7a) and the Mouse Kidney dataset (Supplementary Fig.8a), the segmentation results (red contours) produced by AMM-Seg align closely with the Ground Truth, particularly in regions with intricate edge details and complex morphology. Overall, AMM-Seg not only accurately segments the inner cristae structure but also minimizes errors and prevents over-segmentation. In contrast, other methods exhibit varying levels of blurriness or false positive. In the Human Myocardium dataset (Supplementary Fig.7b), we analyze the F1 scores across three patient groups: Patient#1, Patient#2, and Patient#3. AMM-Seg consistently achieves the highest F1 scores for all patients, clearly outperforming other baseline methods. This demonstrates its superior robustness and accuracy in myocardial tissue segmentation. On the Mouse Kidney dataset (Supplementary Fig.8b), we further compare the methods using IoU, MCC, and F1 score. Compared to F1 alone, IoU and MCC provide complementary views that better reflect performance on complex and fine-grained cristae. AMM-Seg outperforms all other methods across these metrics and shows the lowest performance variance, indicating both high accuracy and strong stability. In contrast, other methods display noticeable fluctuations, failing to balance precision and consistency.

The inner cristae typically exhibit intricate and fine texture features in electron microscope images, presenting challenges for precise segmentation. AMM-Seg employs a dual-channel strategy that extracts augmented and textured images to enhance structural features. Additionally, an adaptive fusion strategy dynamically optimizes the selection of structural features. Furthermore, a multi-domain adaptive strategy is applied to improve its cross-domain capabilities. In summary, these results demonstrate that AMM-Seg consistently outperforms existing methods in segmenting both the mitochondrial outer membrane and the inner cristae, offering superior accuracy, robustness, and cross-domain adaptability.

### Comparison of z-direction segmentation continuity for healthy mitochondria

Original data are acquired by scanning electron microscopy imaging, which exhibits a high degree of continuity between adjacent slices. Therefore, when using a neural network model to segment mitochondria, it is essential not only to accurately segment the inner cristae and outer membranes of mitochondria but also to ensure the continuity of segmentation results between adjacent slices. We evaluate the performance of AMM-Seg on two representative biological datasets: Human Myocardium and Mouse Kidney, focusing on healthy mitochondria for consistency and clarity in structural analysis. Fig.5a and Supplementary Fig.9a present 3D visualizations comparing AMM-Seg with two suboptimal baselines, DA-ISC and MEDIAR. In both cases, xz-plane views highlight inter-slice continuity. AMM-Seg demonstrates greater structural consistency and integrity across both datasets. In regions marked by red circles, it better preserves mitochondrial continuity, while DA-ISC and MEDIAR exhibit segmentation breaks and morphological distortions. These results confirm that AMM-Seg effectively preserves segmentation continuity across adjacent slices, even when handling mitochondria with diverse structural characteristics.

**Fig. 5.**
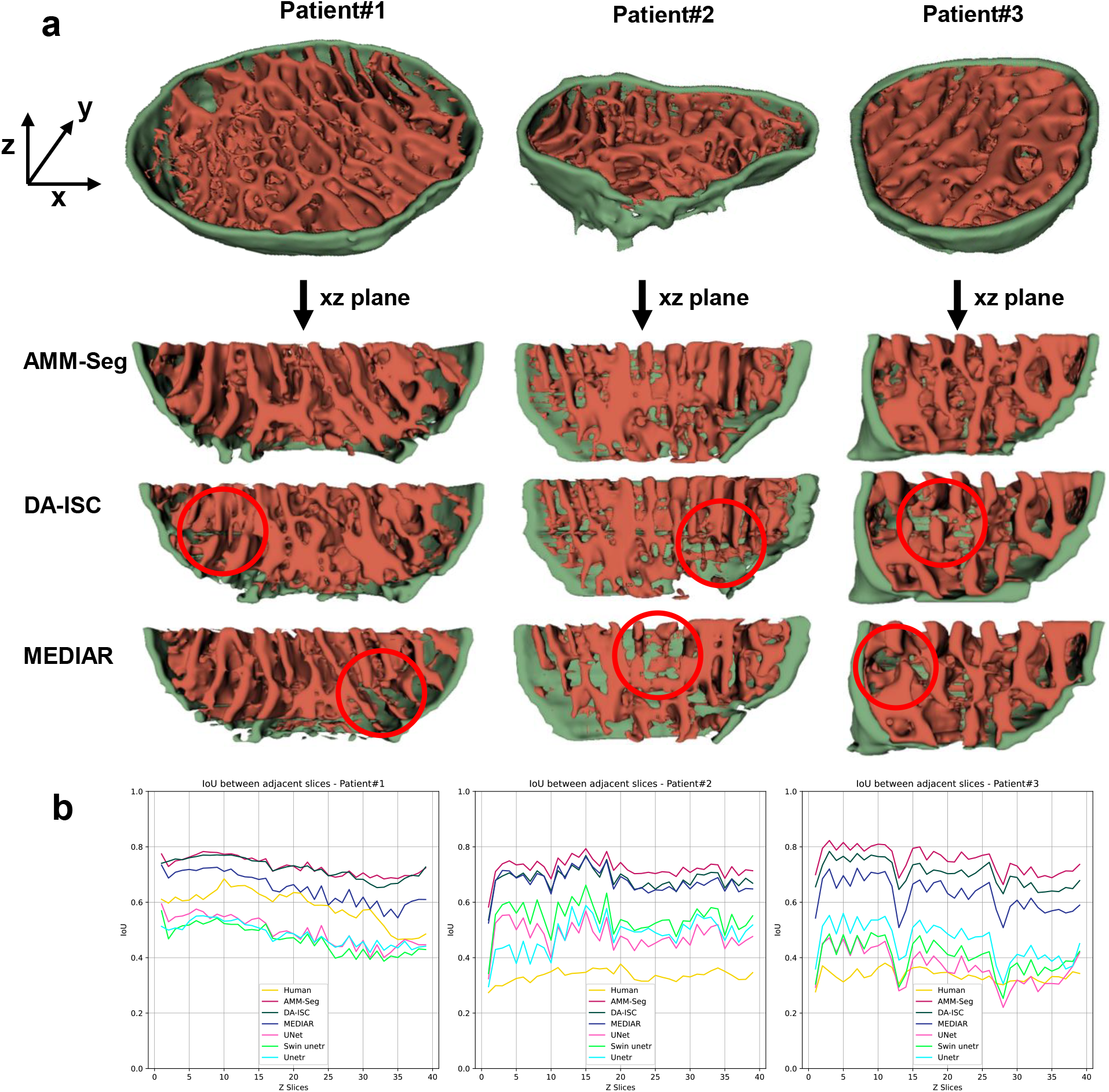
Visual and quantitative comparison of segmentation continuity in the z-direction on the Human Myocardium dataset. **a,** The mitochondrial segmentation results of the three human myocardium patient datasets are visualized through 3D reconstruction using different segmentation methods (AMM-Seg, DA-ISC, MEDIAR). The xz-plane of the 3D reconstruction is extracted to intuitively compare the differences in 3D reconstruction continuity across methods. The red circles highlight areas with poor segmentation continuity observed in DA-ISC and MEDIAR, while AMM-Seg demonstrates more consistent and accurate results. **b,** IoU performance evaluation between adjacent slices. The continuity of 2D image segmentation results is evaluated between adjacent slices for AMM-Seg, six other methods, and human annotation. The IoU curves show that AMM-Seg consistently outperforms DA-ISC, MEDIAR, and other methods across three human myocardium patient datasets, maintaining higher IoU values between adjacent slices. These results suggest that AMM-Seg effectively ensures segmentation consistency in the z-direction through a robust continuity strategy, enabling accurate 3D mitochondrial reconstructions.

After verifying the model’s ability to preserve mitochondrial continuity in 3D visualizations, we further assess its performance by quantitatively evaluating inter-slice consistency. Fig.5b (Human Myocardium dataset) and Supplementary Fig.9b (Mouse Kidney dataset) show the IoU trends across 40 adjacent slices (each 800 × 800 pixels) for AMM-Seg and six other segmentation methods (human means manual annotation). The results indicate that AMM-Seg achieves high stability and continuity in IoU values across slices, with minimal fluctuations, suggesting that its segmentation is more consistent between slices. This advantage is particularly evident in the results for Patient#2 and Patient#3, where AMM-Seg consistently maintains high IoU values that change smoothly along the slice sequence. In contrast, methods such as DA-ISC and MEDIAR exhibit notable drops and fluctuations in IoU across certain slices, reflecting poor inter-slice consistency that may lead to discontinuities and morphological distortions in the final segmentation output.

Overall, the AMM-Seg method is significantly better than other methods in terms of 3D structure preservation of segmentation results and continuity between slices. These experimental results show that AMM-Seg can provide more accurate and stable segmentation results, providing a reliable basis for subsequent 3D reconstruction and quantitative analysis.

### Quantitative analysis of mitochondria in healthy and damaged states

For quantitative analysis, a total of 240 mitochondria are randomly selected, including 60 healthy and 20 damaged mitochondria from each of three human myocardium patient datasets. For each mitochondrion, we assess key structural information: the surface area of the outer membrane, the surface area of the inner cristae, the solid volume (including the volume occupied by the outer membrane and inner cristae), and the total volume (the overall volume of the entire mitochondrion). We then calculate three specific ratios for further analysis: the ratio of inner cristae surface area to total volume, represented as S_inner_/V_total_(Cristae Density^41^), the ratio of inner cristae surface area to outer membrane surface area, represented as S_inner_/S_outer_ and the ratio of solid volume to total volume, represented as V_solid_/V_total_. We obtain an 800 × 800 × 400 voxel structure segmentation of mitochondria using the AMM-seg method. Given that the original image resolution is 5 × 5 × 10 nm^3^ per voxel, we downsample the segmentation in the xy plane to produce 400 × 400 × 400 voxel block, resulting in a uniform pixel size of 10 × 10 × 10 nm^3^. This adjustment provides a consistent scale for real-space representation, facilitating clearer 3D visualization (See Supplementary videos for details).

In Fig.6a, we highlight one healthy mitochondrion in green and one damaged mitochondrion in red, with other mitochondria shown in yellow. The images reveal significant structural differences between healthy and damaged mitochondria: healthy mitochondria exhibit a relatively organized arrangement of inner cristae, while damaged mitochondria show disrupted structure and a disordered cristae arrangement, underscoring the structural changes associated with mitochondrial damage. Secondly, the quantitative statistical results for individual mitochondria are shown in the bottom of Fig.6b.

**Fig. 6.**
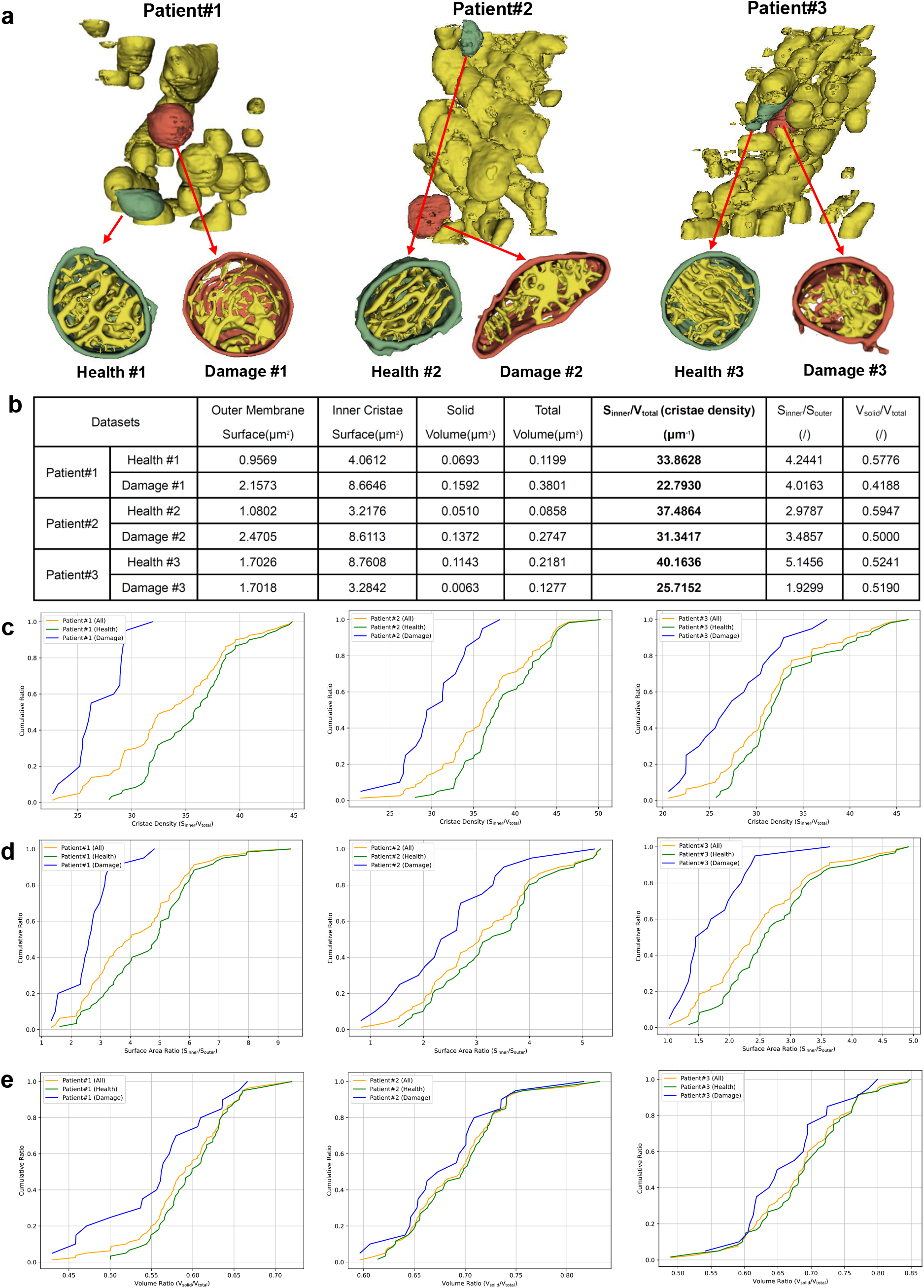
3D visualization and quantitative analysis of healthy and damaged mitochondria in the Human Myocardium dataset. **a,** 3D visualization of segmentation results from three human myocardium patient datasets. In each dataset, one healthy mitochondrion (green) and one damaged mitochondrion (red) are selected. **b,** Quantitative measurements for each selected mitochondrion, including outer membrane surface area, inner cristae surface area, solid volume, total volume, and calculated ratios (S_inner_/V_total_, S_inner_/S_outer_, V_solid_/V_total_). **c,** Cumulative distribution plots of cristae density (S_inner_/V_total_) for each dataset. **d,** Cumulative distribution plots of the surface area ratio (S_inner_/S_outer_) for each dataset. **e,** Cumulative distribution plots of the volume ratio (V_solid_/V_total_) for each dataset. The blue, green, and orange lines in panels **c, d**, and **e** represent damaged, healthy, and all mitochondria, respectively. The three cumulative distribution plots demonstrate that the cumulative distributions of cristae density, surface area ratio, and volume ratio are lower for damaged mitochondria compared to healthy mitochondria, with the most notable difference observed in cristae density.

From these results, the S_inner_/V_total_ (cristae density) values for healthy and damaged mitochondria reveal significant differences between Patient#1 and Patient#3, while the S_inner_/S_outer_ values demonstrate distinct differences between healthy and damaged mitochondria in Patient#3. Thus, a single statistical result does not provide a definitive distinction between healthy and damaged mitochondria. The results show that the S_inner_/V_total_ ratios for Health#1, Health#2, and Health#3 are significantly higher than those for Damage#1, Damage#2, and Damage#3. Combined with 3D visualization, it is evident that the more severe the damage to the inner cristae structure in a mitochondrion, the smaller this S_inner_/V_total_ ratio becomes. This suggests that S_inner_/V_total_ ratio effectively reflects the cristae density of a mitochondrion, supporting the conclusions of Adams et al.^42^. In contrast, the other two ratios—S_inner_/S_outer_ and V_solid_/V_total_—lack consistent trends across datasets and are less effective in distinguishing between healthy and damaged mitochondria. This is primarily because they do not directly capture the architectural complexity of the inner membrane system. The former is sensitive to variations in overall mitochondrial shape, while the latter reflects general structural occupancy without regard to membrane organization. In comparison, S_inner_/V_total_ quantifies the amount of inner membrane surface per unit volume, serving as a proxy for cristae elaboration and thus mitochondrial bioenergetic potential. This makes it a more sensitive and structurally meaningful indicator of mitochondrial health. To further validate the applicability of our analysis, we apply the same quantitative evaluation to the Mouse Kidney dataset, which contains only healthy mitochondria (Supplementary Fig.10).

To achieve a more comprehensive analysis of the differences between healthy and damaged mitochondria, we increase our sample size in three human myocardium patient datasets. Each dataset includes 20 damaged and 60 healthy mitochondria, resulting in a total of 240 mitochondria. We plot the cumulative distribution curves of healthy mitochondria, damaged mitochondria and all mitochondria on the three human myocardium patient datasets, as shown in Fig.6c, Fig.6d and Fig.6e. From the three types of cumulative distribution curves, it is evident that for a given cumulative ratio, the values corresponding to healthy mitochondria are generally higher than damaged mitochondria. This distinction is particularly pronounced in cristae density, where the separation between the distributions of healthy and damaged mitochondria is the most significant, highlighting its effectiveness as a key indicator of mitochondrial health. On the other hand, the volume ratio demonstrates less separation between healthy and damaged mitochondria, indicating that while volume changes may occur during mitochondrial damage, they are less reliable for distinguishing health states. Furthermore, box plots are created with each point representing an individual mitochondrion (Supplementary Fig.11). Additionally, we present statistical data on healthy and damaged mitochondria from the three human myocardium patient datasets, each comprising 20 damaged and 60 healthy mitochondria. The analysis includes the maximum, minimum, median, mean, and standard deviation for each property (Supplementary Table 1-3). Overall, these findings emphasize the importance of integrating multiple structural metrics to achieve a more robust and comprehensive evaluation of mitochondrial integrity. To further facilitate the analysis and visualization of these results, we have developed an intuitive and user-friendly interface (Supplementary Fig.12). This interface supports data import, mitochondrial structure segmentation and the preview and analysis of segmentation results, enabling a streamlined and efficient workflow for researchers.

## Discussion

In this study, we developed MitoStructSeg, a fully automated framework designed to overcome key challenges in both mitochondrial structure segmentation and quantitative analysis. The MitoStructSeg integrates two core modules: the mitochondrial structure segmentation module and the mitochondrial quantitative analysis module. The segmentation module employs the Adaptive Multi-Domain Mitochondrial Segmentation (AMM-Seg) method for accurate segmentation of mitochondrial structures, while the quantitative analysis module provides detailed metrics that are essential for understanding mitochondrial morphology and function.

AMM-Seg demonstrates strong generalization at two levels: across individuals within the same dataset type, and across datasets with distinct mitochondrial structural characteristics. For individual-level generalization (Fig.3),a model trained on one subject generalizes well to other subjects within the same dataset, even with limited annotations. In more challenging settings involving substantial morphological variation (Supplementary Fig.2), the model adapts effectively across datasets and achieves accurate segmentation with only a small amount of labeled data from the target. Building on this segmentation capability, the quantitative analysis module in MitoStructSeg enables comprehensive and reliable measurement of mitochondrial structural features, including surface area, volume, and cristae density. This supports in-depth biological analysis and provides valuable insights into mitochondrial morphology and function (Fig.6).

Theoretically, MitoStructSeg advances the field by showcasing the synergy between mitochondrial structure segmentation and quantitative analysis in biomedical image processing. The AMM-Seg model offers novel insights into the segmentation of complex mitochondrial structures, while the quantitative analysis module delivers precise metrics for investigating mitochondrial morphology and function, thereby enhancing our understanding of the intricate relationship between structure and function. Practically, the MitoStructSeg has the potential to significantly impact biomedical research, particularly in the study of mitochondrial dysfunction and related diseases, where both structural and functional data are critical.

While MitoStructSeg represents a significant advancement, it has limitations that should be addressed in future research. Future work should focus on enhancing the model’s self-learning capabilities, particularly by incorporating large-scale unlabeled datasets for training. Additionally, improving segmentation accuracy for mitochondrial structures by integrating more advanced feature extraction techniques and exploring the use of pre-trained models tailored to specific domains will be essential to broaden its applicability.

In summary, MitoStructSeg makes a significant contribution to mitochondrial structure analysis by delivering superior segmentation accuracy and providing comprehensive quantitative analysis. Its robust performance across diverse datasets and ability to handle unlabeled data establish it as a powerful tool for mitochondrial research. By addressing both structural segmentation and quantitative analysis, this study lays the groundwork for future advancements in image analysis technologies and deep learning applications in biomedical research on mitochondrial dysfunction.

## Methods

### Section 1: Dataset Acquisition

#### Human Myocardium Dataset^43^

In this study, we analyze the myocardial data of three male patients who underwent myocardial biopsy after COVID-19 infection to understand the health of the mitochondrial structure in their myocardium. Patient#1, 30 years old, underwent myocardial biopsy 37 days after infection and is diagnosed with viral myocarditis, sudden cardiac death, and ventricular fibrillation. Patient#2, 44 years old, underwent myocardial biopsy 58 days after infection and is diagnosed with viral myocarditis. Patient#3, 70 years old, underwent myocardial biopsy 76 days after infection and is diagnosed with viral myocarditis, atrial fibrillation, and personal history of cerebral infraction. By selecting these three patients, we obtain a rich data sample with a wide range of time spans and disease diversity, which provides important evidence for in-depth research on the impact of the disease on the myocardium and mitochondria.

The myocardial tissue samples are sourced from the three patients mentioned above, and the process for obtaining the data required for AI model training is as follows. Myocardial biopsy samples from COVID-19 patients are sectioned into 1 × 1 mm^2^ blocks and pre-fixed in a pH 7.2 solution containing 2% paraformaldehyde and 2.5% glutaraldehyde for 12 hours. The samples then underwent osmiumthiocarbohydrazide-osmium (OTO) staining. They are fixed in 1% osmium tetroxide with potassium ferrocyanide for 1 hour on ice, washed with phosphate buffer and ddH2O, treated with 1% thiocarbohydrazide, and post-fixed in 1% osmium tetroxide. Following staining with 1% uranium acetate and 0.66% lead citrate, the samples are dehydrated through a graded ethanol and acetone series, infiltrated with Epon 812 resin, and polymerized at 60°C for 48-72 hours. The resin blocks are observed under a dual-beam scanning electron microscope (Aquilos Cryo-FIB, Thermo Fisher Scientific). Each serial face is imaged using the T1 (in-lens detector) backscatter mode with an acceleration voltage of 2.0 keV and a current of 0.1 nA. Sequential milling is performed using a gallium ion beam, with a Z-axis height of 10 nm for each milling.

According to the above process, we obtain the data from three patients. The data sizes are 6144 × 4096 × 800 voxels for Patient#1, 6144 × 4096 × 800 voxels for Patient#2, and 6500 × 5200 × 800 voxels for Patient#3. The pixel size is 5 × 5 nm^2^ and the slice interval is 10 nm. Initially, we crop two regions (800 × 800 × 100 voxels) from the data of Patient#1 and manually annotated them to create the source domain dataset (with labels). Subsequently, we crop a region (800 × 800 × 200 voxels) from the data of Patient#1, Patient#2, and Patient#3 respectively to form the target domain dataset (without labels). Additionally, we crop a region (800 × 800 × 40 voxels) from the data of these three patients to create the validation dataset for evaluating the segmentation accuracy and generalization performance of AMM-Seg. The Human Myocardium Dataset is available at https://github.com/xiaohuawan/MitoStructSeg.

#### Mouse Kidney Dataset

We also conduct mitochondrial structure segmentation experiments on the Mouse Kidney dataset. This publicly available dataset consists of kidney tissue from a wild-type, healthy, 8-week-old mouse obtained from Jackson Laboratory. The original data volume has dimensions of 12287 × 7976 × 22199 voxels, with an isotropic voxel size of 8 × 8 × 8 nm^3^. A region of 800 × 800 pixels is cropped and 50 slices are manually annotated to serve as part of the source domain dataset (with labels), together with the 200 labeled slices from the Human Myocardium Dataset. Additionally, another region of size 800 × 800 × 200 voxels is extracted as the target domain dataset (without labels). Finally, 40 slices are selected from this dataset to evaluate the segmentation accuracy and generalization performance of AMM-Seg. The Mouse Kidney dataset is available at https://doi.org/10.25378/janelia.16913035.v1.

### Section 2: AMM-Seg Model And Training

The AMM-Seg model (Supplementary Fig.1) addresses mitochondrial segmentation challenges in noisy and complex imaging environments by leveraging a domain adaptation framework that minimizes labeling costs and improves model robustness. This framework uses both a labeled source domain and an unlabeled target domain, allowing for adaptive learning in the target domain and reducing dependency on manually labeled data. The core architecture of AMM-Seg is a dual-channel structure that extracts distinct image features, which are separated into augmented and textured features. An adaptive fusion module then integrates these features, enhancing segmentation accuracy for complex mitochondrial structure like cristae by preserving crucial texture details. To guide the model training across domains, we use specific loss functions, including a source domain segmentation loss, a target domain adaptation loss, and a domain classification loss. The following sections describe each module of the AMM-Seg model in detail.

#### Dual-channel features Module

To enhance generalization, reduce overfitting, and improve segmentation accuracy for fine mitochondrial structure such as cristae and membranes, we apply tailored data augmentation and texture extraction techniques to each domain. In the source domain, weak geometric augmentation (𝒜 *_g_*) prevents overfitting by applying transformations such as resizing, cropping, and flipping to generate augmented images, represented as 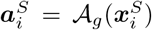. For the target domain, random intensity-based augmentation (𝒜 *_r_*) is used to enhance robustness by varying contrast and brightness, resulting in 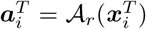. Additionally, to capture essential texture details, we preprocess the images using a Gaussian blur to reduce noise and highlight significant structural patterns. Following this, a Canny edge detection filter is applied to extract image texture. This process generates texture-enhanced images, represented as 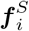 and 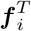 for the source and target domains, respectively. These preprocessing steps provide robust feature representations across diverse imaging conditions, facilitating accurate segmentation of intricate mitochondrial structure.

#### Adaptive fusion Module

For feature encoding, we employ the Cross-Slice Fusion (CSF) module to combine adjacent slices during downsampling, which retains continuity across the 2D mitochondrial slices. This operation is defined as:

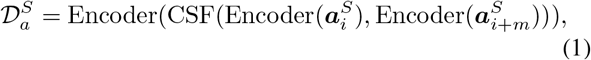

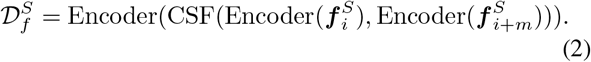

where m=1, which means fusing adjacent images. And Encoder(·) represents the downsampling operation. CSF is applied to both augmented and texture-enhanced features, creating unified representations across adjacent slices. Similarly, for the target domain, we obtain 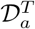 and 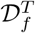.

To dynamically integrate encoding features, we design a channel adaptive fusion module inspired by the self-attention mechanism^44^. This module computes query, key, and value matrices ***Q****^S^*, ***K****^S^*, and ***V****^S^* based on the channel attention representations of augmented and textured features, denoted as 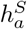 and 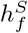, as follows:

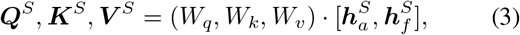

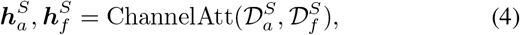

where *W_q_, W_k_*, and *W_v_* are learnable matrices. The fused representation 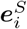 is then obtained by combining the encoded features with attention weights ***w_a_***and ***w_f_***as follows:

Similarly, we can obtain the encoding result 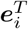 in the target domain. It is worth noting that the encoder of augmented and texture image features adopts a weight-sharing strategy to reduce model complexity and improve computational efficiency. To effectively process segmentation and inter-slice variations, we employ a dual-decoder design. One decoder is dedicated to segmentation outputs (Equations (7) and (8)), while the other is designed to capture inter-slice variations (Equation (9). The Slice Decomposition (SD) module further separates adjacent slice features, defined as:

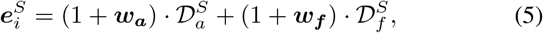

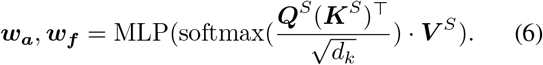

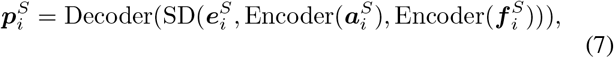

where Decoder and Up_diff_ are upsampling operations, but they differ in network structure. 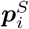 and 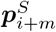 is the segmentation result and 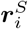 is the feature of image variation. Similarity allows us to obtain 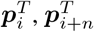, and 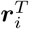 for the target domain.

##### Loss Function

To optimize segmentation quality and enable effective cross-domain adaptability, we design three types of loss functions: source domain loss, target domain loss, and domain classification loss.

The source domain loss ensures accurate segmentation in the labeled source domain through cross-entropy loss, addressing both segmentation performance and inter-slice continuity:

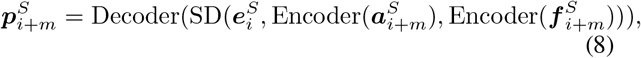

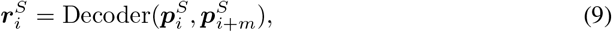

where *CE*(·) represents cross-entropy loss, and XOR(·) captures label differences between adjacent slices. The target domain loss addresses the lack of labels in the target domain by generating pseudo-labels 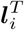 and 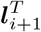 from model predictions. Cross-entropy loss is computed for the target domain as follows:

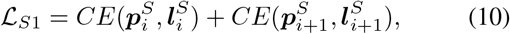

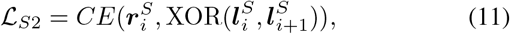

To achieve domain-invariant feature extraction, we use a domain classification loss with a gradient reversal layer (GRL)^45^, guiding a domain classifier to differentiate between source and target domain data:

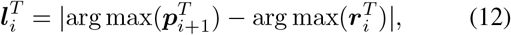

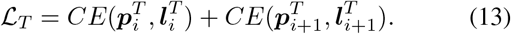

where *G_d_* is domain classification function, 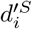 and 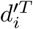 are the domain labels.

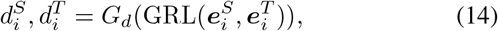

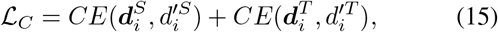

The total loss function, ℒ, combines these three loss components, with *λ*_1_ and *λ*_2_ controlling the contributions of target domain loss and domain classification loss, respectively:

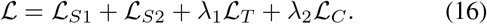

#### Training details

For model training, we adopt a transfer learning strategy that addresses both within-dataset and across-dataset generalization. For the Patient#1 dataset (Human Myocardium), the model is trained in a semi-supervised setting using 200 labeled slices and 200 unlabeled slices from the same individual. For the Patient#2 and Patient#3 datasets (Human Myocardium), the model is evaluated on its ability to generalize across individuals with different mitochondrial structural characteristics. Specifically, 200 labeled slices from Patient#1 are used for training, while 200 unlabeled slices from Patient#2 or Patient#3 are used for testing. For the Mouse Kidney dataset, the model is trained to adapt across datasets with distinct mitochondrial morphologies. In this setting, 200 labeled slices from Patient#1 and 50 labeled slices from the Mouse Kidney dataset are used for training, and 200 unlabeled slices from the Mouse Kidney dataset are used for evaluation. The training process begins with an initial learning rate of 0.0005, which is dynamically adjusted throughout the training period using a polynomial decay strategy with a decay power of 0.9. This strategy reduces the learning rate progressively over a total of 10,000 iterations, promoting stable convergence and helping to prevent early overfitting. The optimization is conducted using the Adam optimizer^46^, configured with *β*_1_ = 0.9 and *β*_2_ = 0.99. A batch size of 2 is employed, with a random selection of samples at each iteration. Given this batch size, the model completes a total of 10,000 effective iterations across the dataset. The parameters of the loss function, as specified in Equation (16), are consistently set to *λ*_1_ = 0.1 and *λ*_2_ = 0.1.

To ensure consistency and comparability across experiments, all datasets are trained using identical configuration settings, including optimizer parameters, learning rate schedule, batch size, and loss weights.

### Section 3: Other Supplementary

#### Code avalidability

The AMM-Seg segmentation algorithms are trained using pytorch 2.4.1, and the quantification algorithms are implemented in Python 3.8.19 using the libraries: albumentations 1.4.7, numpy 1.24.4, opencv-python 4.9.0.80, scikit-image 0.21.0, scipy 1.10.1, and pandas 2.0.3. These algorithms are accelerated using the GPUs (NVIDIA A100-PCIE-40GB) on CentOS Linux 7. The GUI is built using React 18.3.1 and Vite 5.2.10, integrated with Ant Design 5.19.3 and Material UI component libraries to provide a modern user interface and responsive layout. The project is developed using Node.js 18.17.1 and JavaScript, with all dependencies managed through npm 9.6.7. The data visualization section incorporates Chart.js 4.4.3, Echarts 5.5.1, and ApexCharts 3.46.2, supporting dynamic display of various types of charts. Additional details, source code, and documentation are available at https://github.com/xiaohuawan/MitoStructSeg.

#### 3D visualization

3D visualization of mitochondrial structure is performed using 3D Slicer version 5.6.2^36^.

## Acknowledgements

This work was supported in part by the National Natural Science Foundation of China (No.32241027, 62472034, 62227807, 92469111, 32241028).

## Author contributions

X.W. and X.Wa. designed the experiments and wrote the paper. X.W., B.C., Z.J., and Y.C. performed the experiments. S.G., Z.L. and F.L. assisted with data collection and preprocessing. X.Wa., Z.L., F.Z. and B.H. secured funding. All have commented on and edited the manuscript.

