## supplementary for "MitoStructSeg: mitochondrial structural complexity resolution via adaptive learning for cross-sample morphometric profiling"

### 1 Supplementary Figures

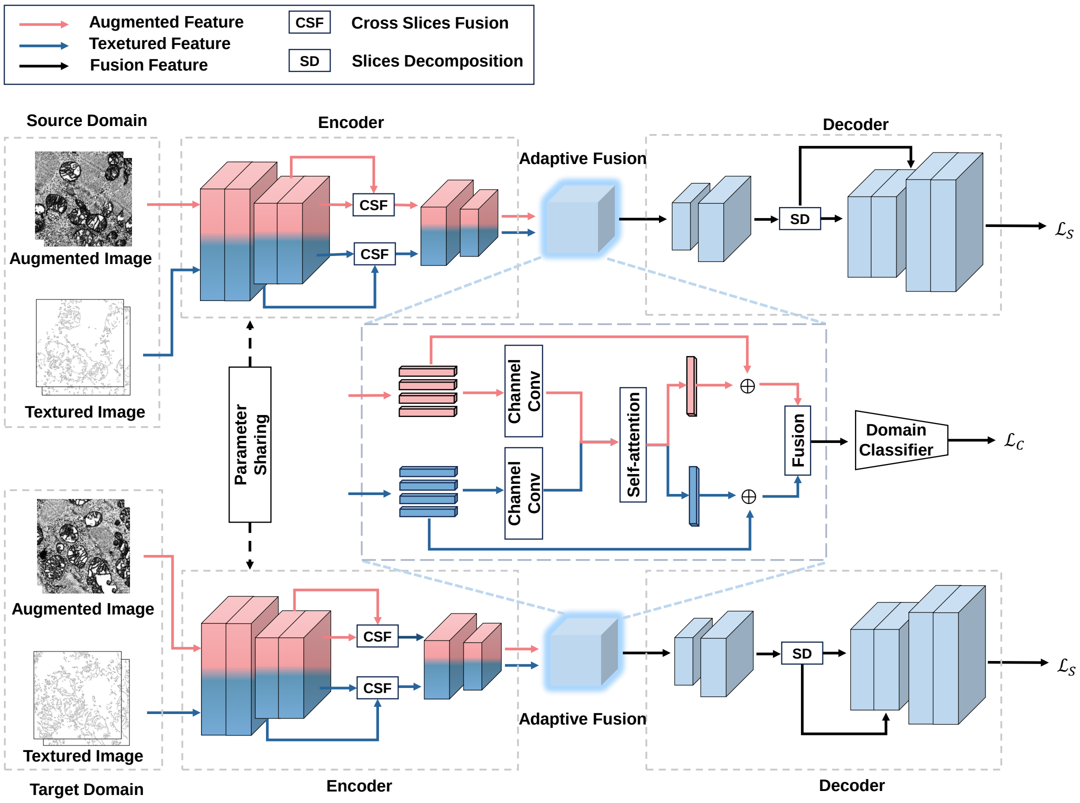


**Supplementary Fig. 1 Summary of the AMM-Seg architecture. Multi-domain adaptation strategy,** AMM-Seg adopts a multi-domain adaptation strategy, using labeled data as the source domain and unlabeled data as the target domain, enhancing domain transferability and overall performance by extracting shared features across domains. **Adaptive fusion strategy,** For in-domain data, a dual-channel strategy extracts augmented image and textured image to strengthen structural feature. Additionally, an adaptive fusion strategy dynamically optimizes the selection of structural features. **Continuity strategy,** Due to the 3D structural characteristics of mitochondria, we fuse the features of adjacent slices through the Cross Slices Fusion (CSF) module and separate them using the Slices Decomposition (SD) module. This process enables the model to learn the continuity features between adjacent slices, further preserving the structural features of mitochondria in the Z-direction.

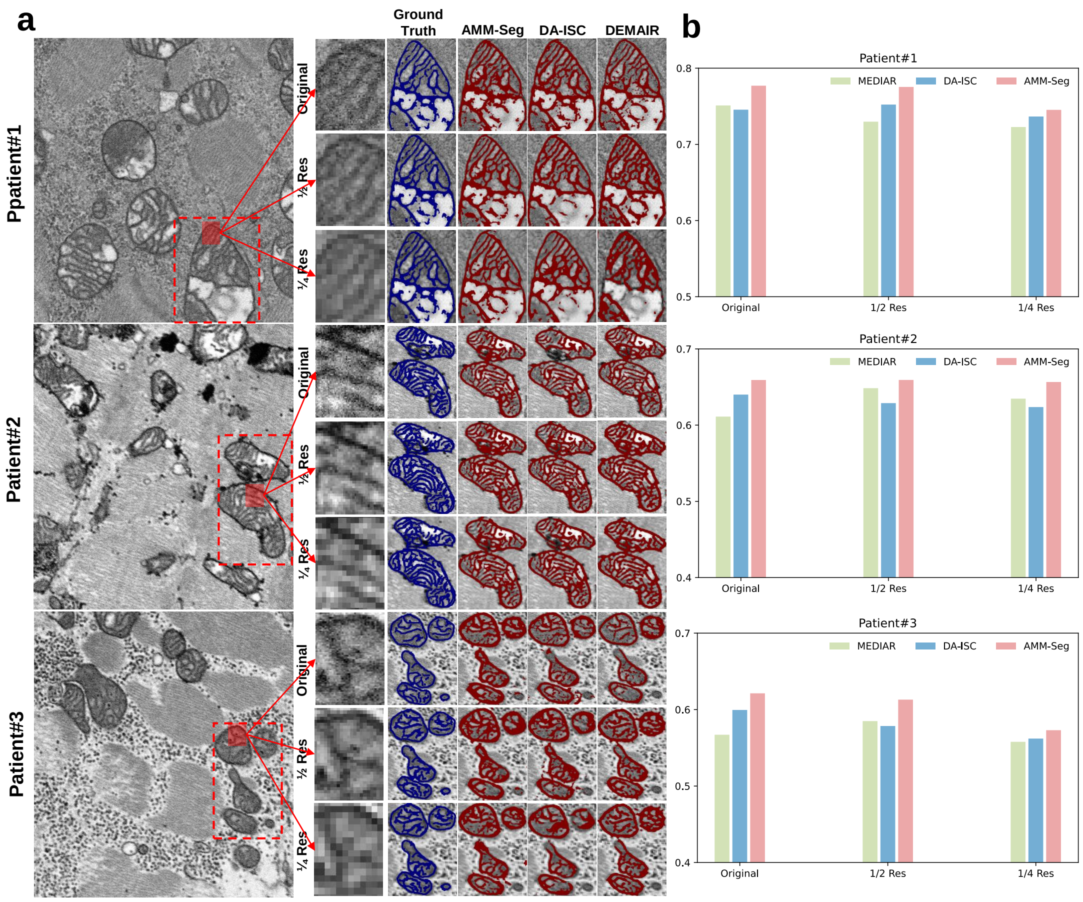


**Supplementary Fig. 2 Comparison of AMM-Seg and suboptimal methods for structure segmentation at different resolutions in the Human Myocardium dataset. a,** The segmentation results of images with different resolutions are shown, namely original resolution, 1/2 original resolution, and 1/4 original resolution. The first column is the enlarged area of the red box in the image, where the difference in image resolution can be seen more clearly. As the resolution decreases, the details of the image gradually blur. Notably, the AMM-Seg method maintains excellent segmentation ability, with its performance showing no significant decline compared to other methods. **b,** The bar chart shows the performance comparison of the three segmentation methods (AMM-Seg, DA-ISC, and MEDIAR) on images with different resolutions. It is evident that the AMM-Seg method achieves the highest segmentation accuracy at all resolutions, maintaining excellent performance at 1/2 and 1/4 of the original resolution, demonstrating its adaptability and robustness to resolution changes.

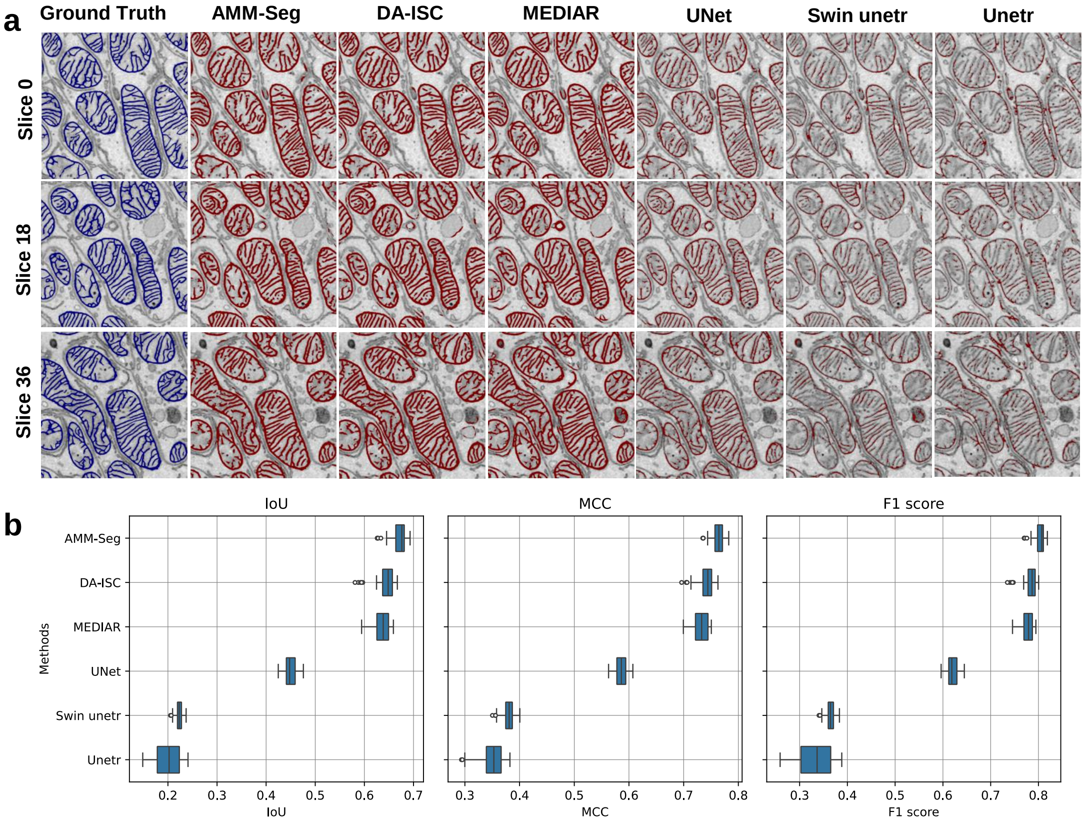


**Supplementary Fig. 3 Comparison of AMM-Seg and mainstream methods for structure segmentation in the Mouse Kidney dataset. a,** The segmentation results of AMM-Seg and five mainstream methods are shown on three representative slices (Slice 0, 18, and 36), which are sampled at an interval of 18 slices from the validation set. The ground truth (blue contours) and predicted results (red contours) are overlaid on the raw EM images. AMM-Seg exhibits higher consistency with the ground truth compared to other methods, especially in capturing fine mitochondrial structures. **b,** Quantitative evaluation is conducted using three standard segmentation metrics: IoU, MCC, and F1 score. AMM-Seg achieves the highest median values with minimal variability, demonstrating both high accuracy and strong robustness. In contrast, UNet, Swin unetr, and Unetr perform poorly, exhibiting low scores and large fluctuations. While DA-ISC and MEDIAR outperform the baseline models to some extent, their performance remains consistently inferior to that of AMM-Seg across all metrics. These results underscore the effectiveness of AMM-Seg in structure segmentation task on the Mouse Kidney dataset.

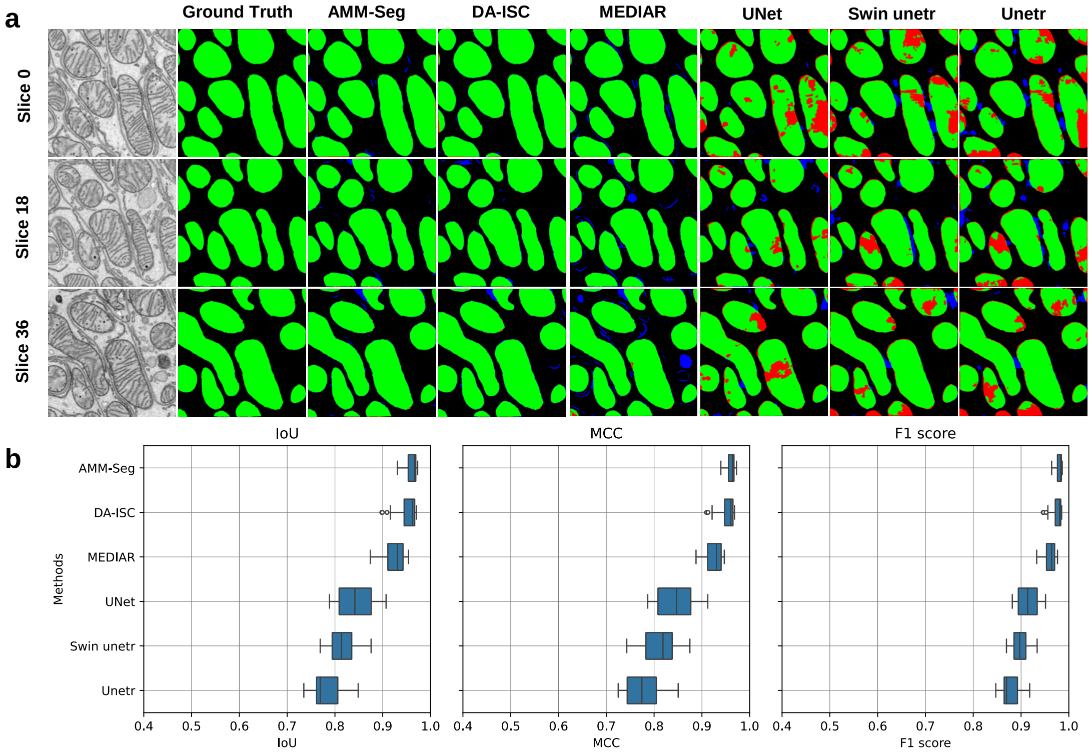


**Supplementary Fig. 4 Comparison of AMM-Seg and mainstream methods for block segmentation in the Mouse Kidney dataset. a,** The results of AMM-Seg, DA-ISC, and MEDIAR are obtained using Flood Fill based on mitochondrial structure segmentation, while UNet, Swin unetr, and Unetr are retrained models for block segmentation task. The segmentation results on three representative slices (Slice 0, 18, and 36) are shown. For each method, the correct part predicted (green), the missing predicted part (red), and the wrong predicted part (blue) are visualized. The results show that AMM-Seg produces predictions more consistent with the Ground Truth, with significantly fewer missing (red) and incorrect (blue) regions compared to other methods. **b,** Quantitative results show that AMM-Seg achieves the highest median IoU, MCC, and F1 score with low variability, indicating strong performance and robustness. While DA-ISC and MEDIAR offer some improvements over baseline models, their performance still falls short of AMM-Seg. These findings underscore AMM-Seg's strength in accurately segmenting complex structures.

---

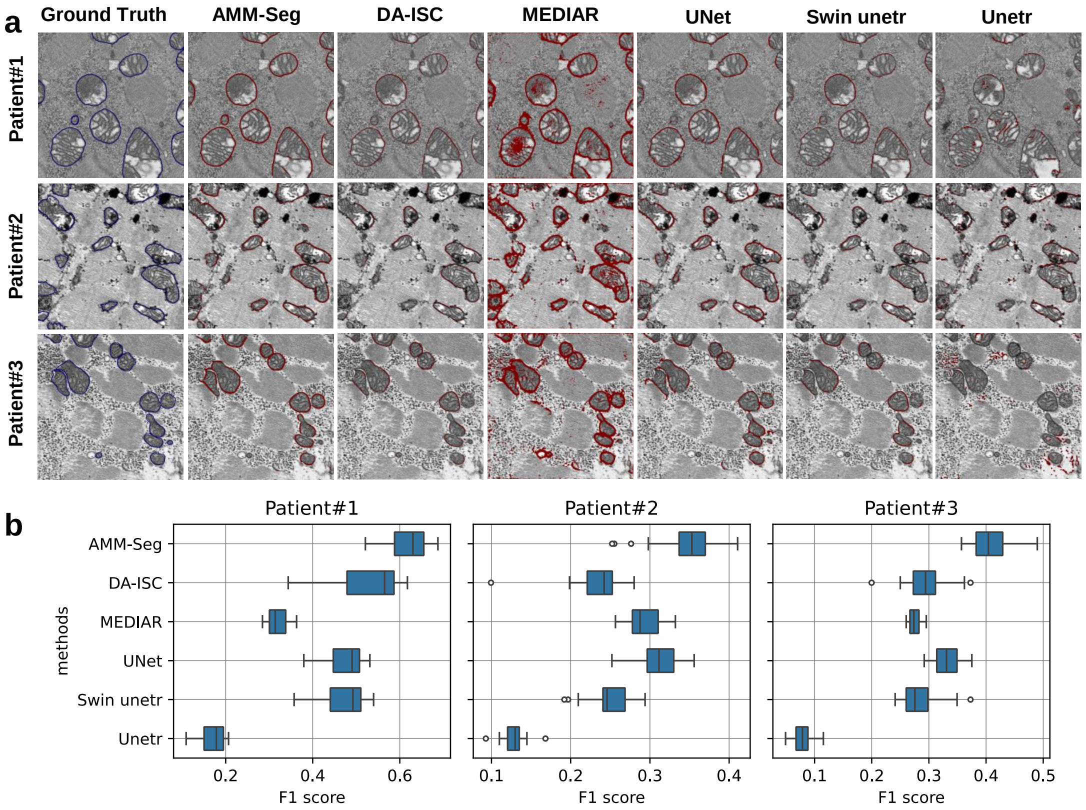


**Supplementary Fig. 5 Comparison of AMM-Seg and mainstream methods for outer membrane segmentation in the Human Myocardium dataset. a,** The AMM-Seg method demonstrates superior segmentation accuracy across three human myocardium patient datasets, with its red contours closely matching the ground truth (blue contours). In contrast, the performance of other methods varies widely, often predicting numerous incorrect regions. For instance, MEDIAR and Unetr tend to introduce substantial noise, resulting in many false positive regions that deviate from the true outer membrane structure. **b,** The quantitative comparison illustrates that AMM-Seg achieves the highest median F1 score, with minimal fluctuations across three human myocardium patient datasets, indicating superior accuracy and robustness. Other methods exhibit both lower median scores and greater fluctuations across datasets, signifying decreased consistency and robustness in segmentation performance. The results indicate that AMM-Seg demonstrates strong capability in outer membrane segmentation task.

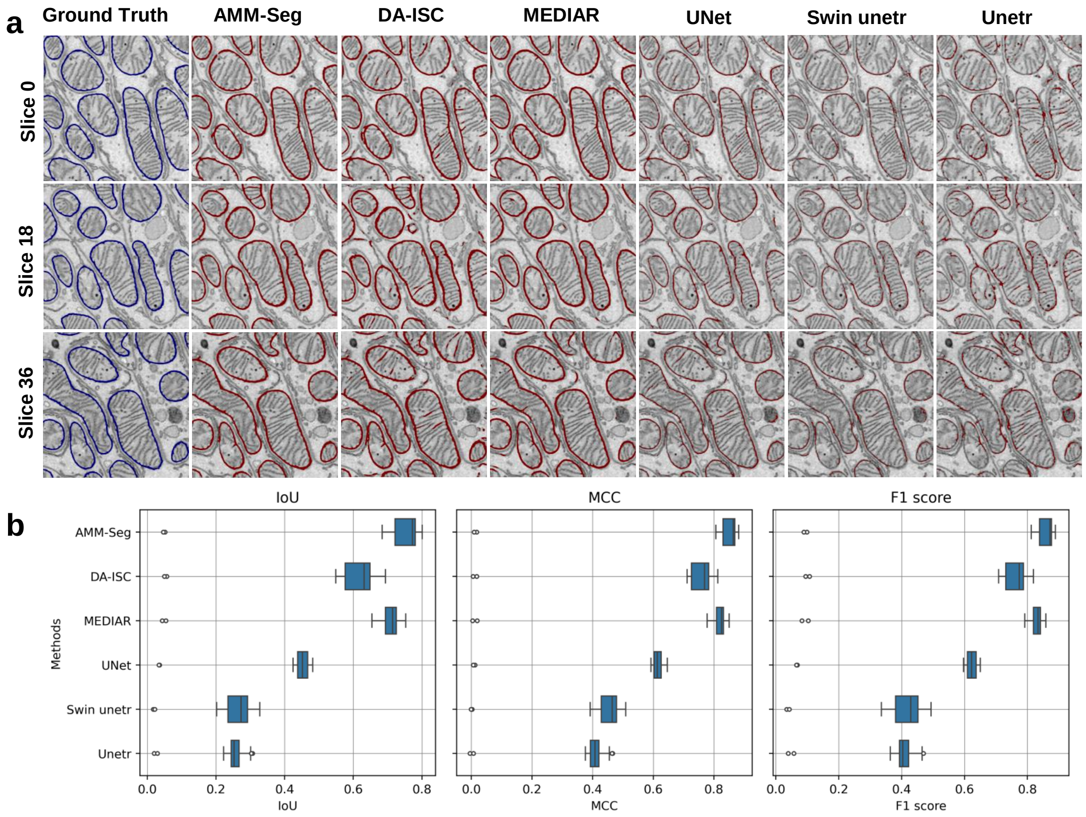


**Supplementary Fig. 6 Comparison of AMM-Seg and mainstream methods for outer membrane segmentation in the Mouse Kidney dataset. a,** The AMM-Seg method demonstrates consistent and high-precision segmentation across three representative image slices (Slice 0, 18, and 36), with red contours closely aligning with the ground truth boundaries (blue contours). In contrast, baseline methods such as DA-ISC, MEDIAR, and UNet often deviate from the true boundaries, with frequent over-segmentation and mismatches. DA-ISC and UNet produce noisier, fragmented results, while Swin unetr and Unetr tend to under-segment, missing key membrane regions. **b,** The quantitative evaluation confirms AMM-Seg's superior performance, achieving the highest median values across all three metrics (IoU, MCC, and F1 score). In contrast, other methods yield lower median scores and wider spread distributions, highlighting their instability and reduced segmentation reliability. These results underscore the effectiveness of AMM-Seg in outer membrane segmentation task on the Mouse Kidney dataset.

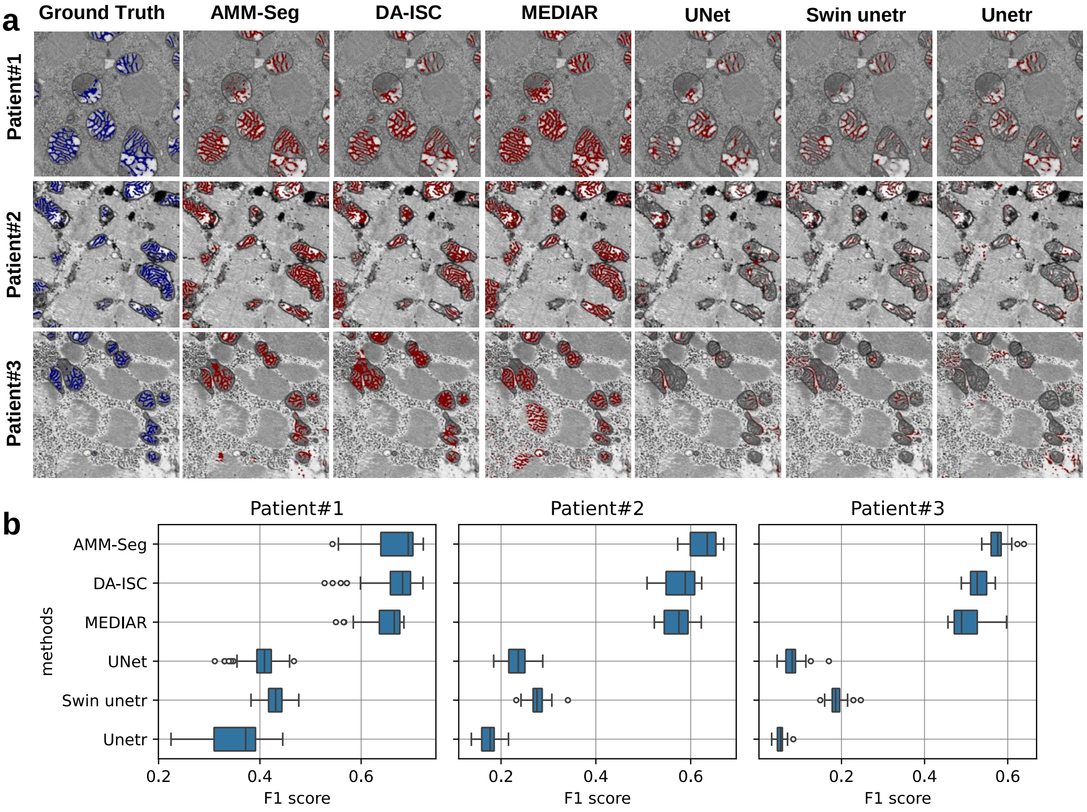


**Supplementary Fig. 7 Comparison of AMM-Seg and mainstream methods for inner cristae segmentation in the Human Myocardium dataset. a,** The AMM-Seg demonstrates superior accuracy compared to other methods, closely matching the ground truth annotations. Visual results indicate that AMM-Seg and DA-ISC perform similarly in capturing cristae structures. MEDIAR, while effective in some areas, tends to mistakenly segment non-cristae structures. The remaining methods—UNet, Swin Unetr, and Unetr—show noticeably lower accuracy, struggling to consistently delineate the cristae. **b,** AMM-Seg achieves the highest median F1 scores, highlighting its robustness and accuracy across all datasets. In comparison, the remaining methods exhibit both lower median F1 scores and broader score distributions, particularly in Patient#2 and Patient#3, indicating significant fluctuations in performance. These results confirm that AMM-Seg offers the most reliable segmentation results among the evaluated methods for mitochondrial cristae segmentation.

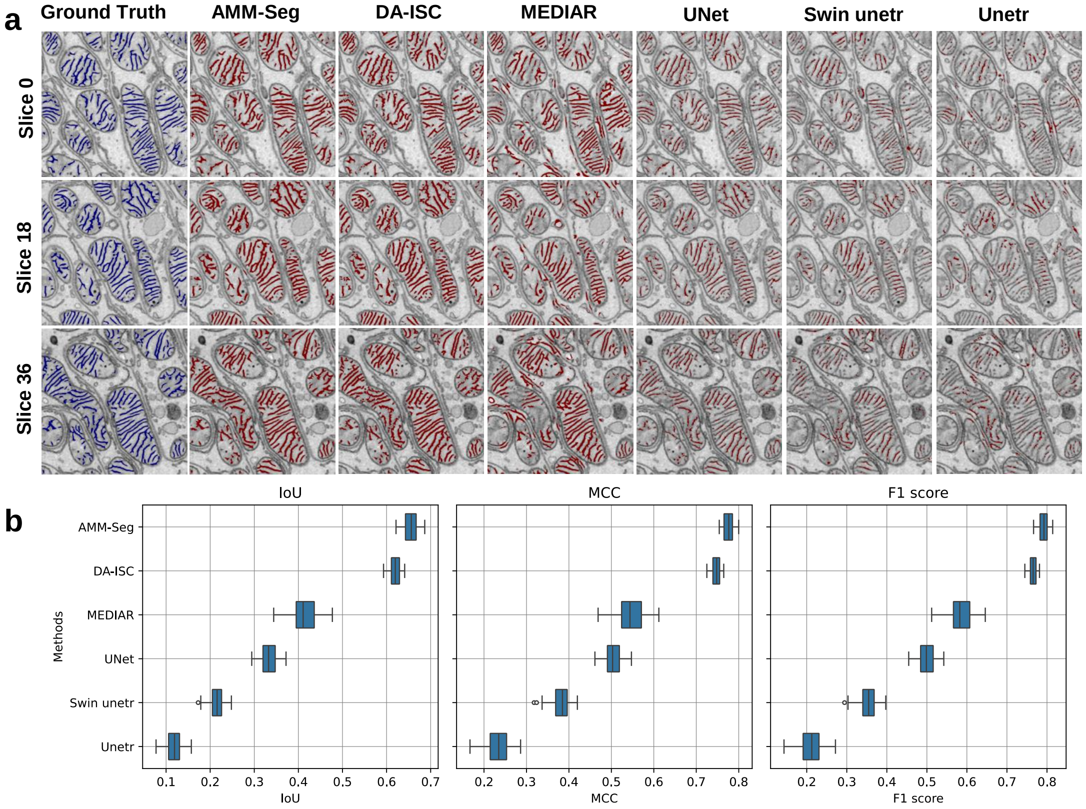


**Supplementary Fig. 8 Comparison of AMM-Seg and mainstream methods for inner cristae segmentation in the Mouse Kidney dataset. a,** The AMM-Seg most accurately captures the fine cristae structure on three representative image slices (Slice 0, 18, and 36), closely matching the ground truth. DA-ISC also delivers competitive performance, producing results that in many regions approximate those of AMM-Seg. MEDIAR shows moderate effectiveness but tends to misclassify non-cristae regions, leading to reduced overall accuracy. In comparison, UNet, Swin unetr, and Unetr consistently underperform, often failing to capture the intricate morphology of cristae and producing fragmented or incomplete segmentations. **b,** AMM-Seg demonstrates superior performance in IoU, MCC, and F1 score, achieving the highest median values and the lowest variability, which reflects its accuracy and robustness. DA-ISC ranks second across all metrics, with performance close to AMM-Seg and relatively low variability. MEDIAR shows moderate performance with less stable segmentation results across slices. In contrast, UNet, Swin Unetr, and Unetr achieve noticeably lower scores with higher variance, indicating reduced reliability and consistency. These results underscore the effectiveness of AMM-Seg in inner cristae segmentation task on the Mouse Kidney dataset.

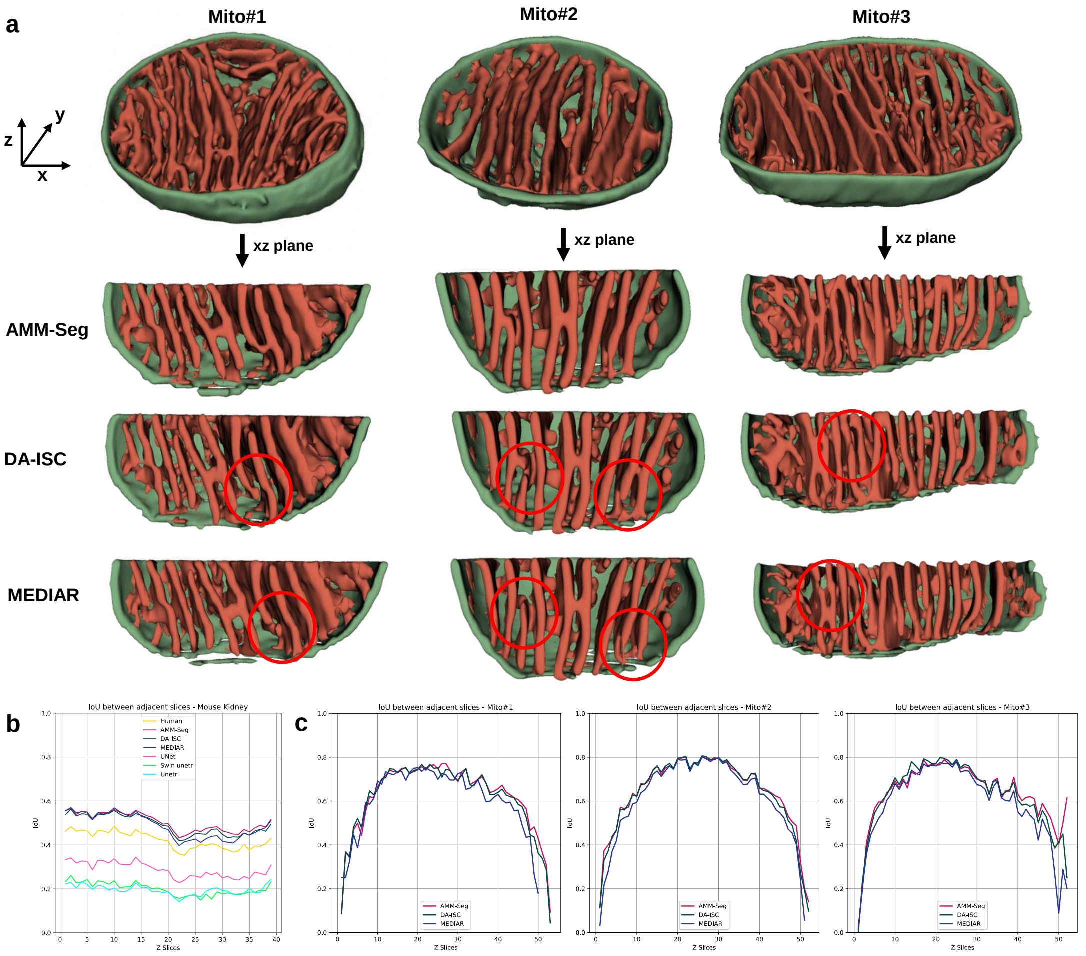


**Supplementary Fig. 9 Visual and quantitative comparison of segmentation continuity in the z-direction on the Mouse Kidney dataset. a,** Three representative mitochondria (Mito#1, Mito#2, and Mito#3) are selected from the Mouse Kidney dataset. Their segmentation results are visualized using xz-plane cross-sections to examine continuity across adjacent slices. Red circles highlight discontinuities or structural inconsistencies in DA-ISC and MEDIAR segmentations. AMM-Seg provides visually smoother and more continuous reconstructions across all three samples. **b,** IoU performance is evaluated between adjacent slices within a 40-slice region (800×800 pixels) selected from the Mouse Kidney dataset. This analysis includes comparisons among AMM-Seg, six other segmentation methods, and human annotation. AMM-Seg consistently achieves the highest IoU values, indicating superior slice-to-slice consistency in this region. **c,** To assess segmentation continuity at the single-mitochondrion level, we compute the IoU between adjacent slices for Mito#1, Mito#2, and Mito#3. AMM-Seg consistently shows smoother and more stable IoU curves than DA-ISC and MEDIAR, aligning with the overall trend observed in panel **b**.

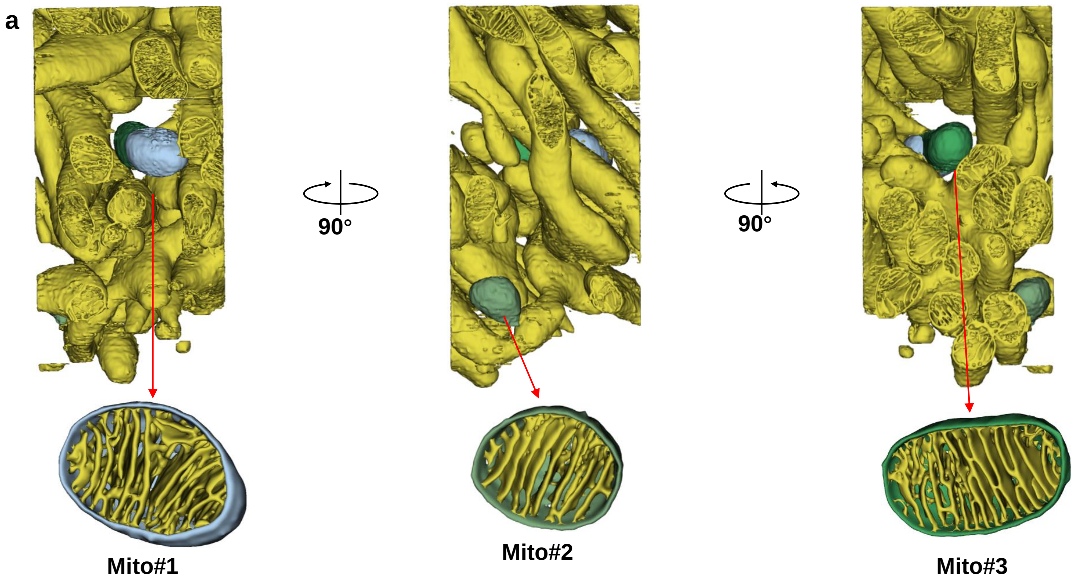


b

| Mito | Outer Membrane Surface(μm²) | Inner Cristae Surface(μm²) | Solid Volume(μm³) | Total Volume(μm³) | $S_{inner}/V_{total}$ (cristae density) (μm⁻¹) | $S_{inner}/S_{outer}$ (/) | $V_{solid}/V_{total}$ (/) |
|---|---|---|---|---|---|---|---|
| Mito#1 | 2.1688 | 11.2219 | 0.3903 | 0.7753 | 14.4743 | 5.1742 | 0.5034 |
| Mito#2 | 1.6242 | 6.5626 | 0.2433 | 0.4764 | 13.7745 | 4.0405 | 0.5107 |
| Mito#3 | 2.3762 | 13.1789 | 0.4406 | 0.8837 | 14.9138 | 5.5463 | 0.4986 |

**Supplementary Fig. 10 3D visualization and quantitative analysis of healthy mitochondria in the Mouse Kidney dataset. a,** 3D visualization of segmentation results for three representative healthy mitochondria (Mito#1, Mito#2, Mito#3) extracted from a single tissue block in the Mouse Kidney dataset. All three views are based on the same tissue block centered on Mito#2, with left and right 90° rotations used to obtain the views of Mito#1 and Mito#3. **b,** Quantitative measurements for the selected mitochondria, including outer membrane surface area, inner cristae surface area, solid volume, total volume, and calculated ratios($S_{inner}/V_{total}$, $S_{inner}/S_{outer}$, $V_{solid}/V_{total}$).

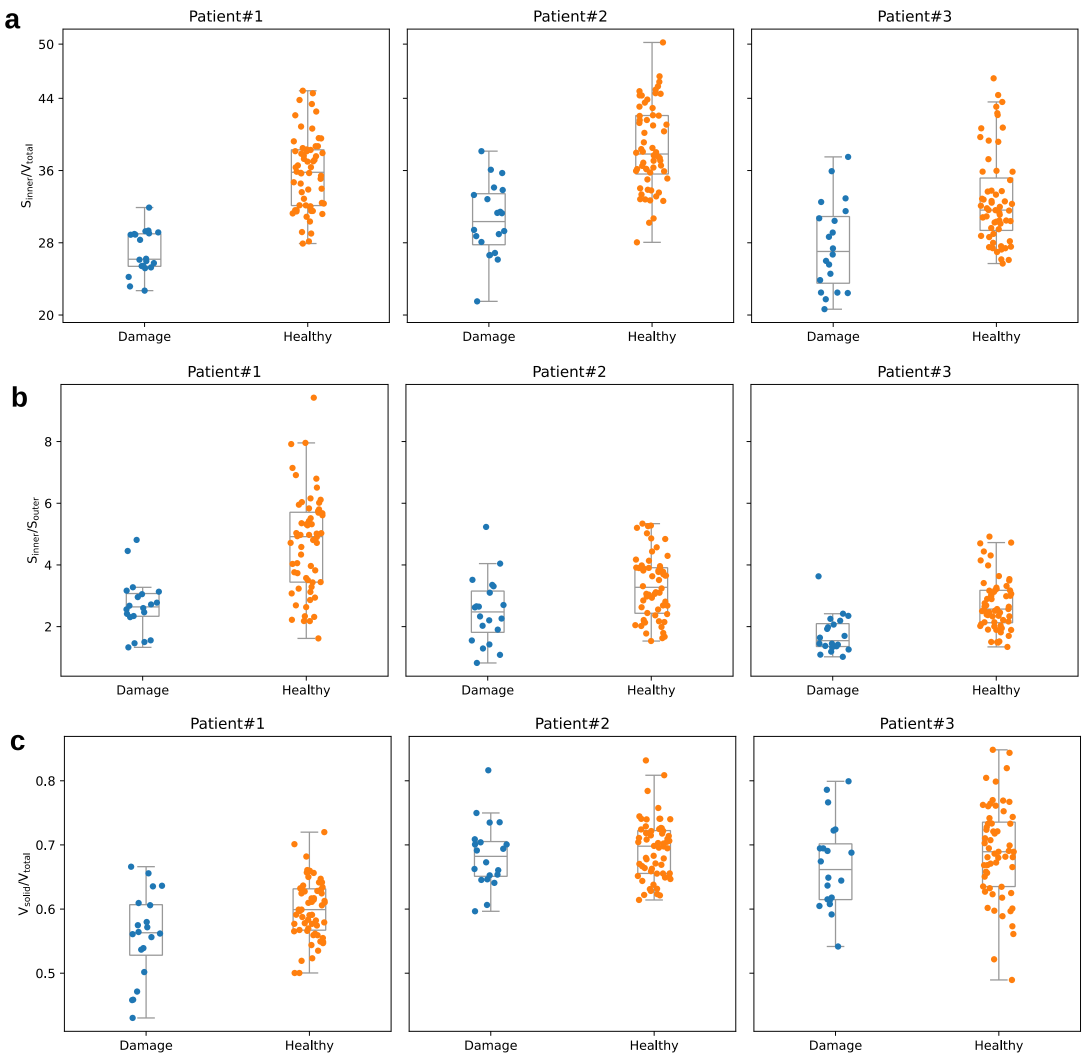


**Supplementary Fig. 11 Quantitative analysis of healthy and damaged mitochondria using box plots in the Human Myocardium dataset. a,** Box plots of the cristae density($S_{inner}/V_{total}$) for three human myocardium patient datasets. **b,** Box plots of the surface area ratio ($S_{inner}/S_{outer}$) for three human myocardium patient datasets. **c,** Box plots of the volume ratio($V_{solid}/V_{total}$) for three human myocardium patient datasets. The blue and orange dots represent damaged and healthy mitochondria, respectively. The three box plots demonstrate that the median values of cristae density, surface area ratio and volume ratio are lower for damaged mitochondria compared to healthy mitochondria, with the most notable difference observed in cristae density.

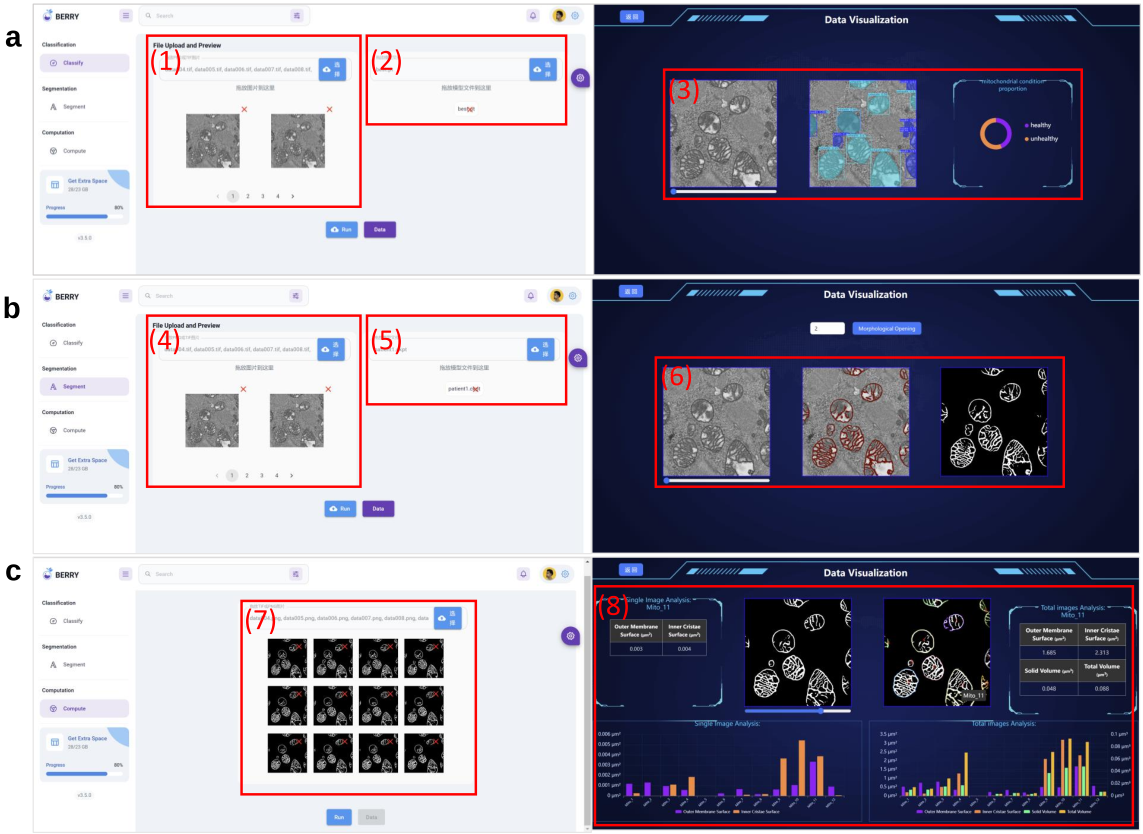


**Supplementary Fig. 12 Overview of the User Interface of the Platform. a,** The mitochondrial classification module categorizes mitochondria into healthy and damaged classes based on user-provided data. (1) Import images for classification. (2) Load pre-trained classification model. (3) Display mitochondrial classification results. **b,** The mitochondrial structure segmentation module performs precise image segmentation to delineate mitochondrial structures. (4) Import images for segmentation. (5) Load pre-trained segmentation model. (6) Display mitochondrial segmentation results. **c,** The mitochondrial quantitative analysis module computes relevant statistical information from the segmentation results and presents it in real time. (7) Import mitochondrial segmentation results. (8) Display statistical information on mitochondria.

#### 2 Supplementary Tables

**Supplementary Table 1** Patient#1 dataset mitochondria properties.

| | | Outer Membrane Surface($\mu m^2$) | Inner Cristae Surface($\mu m^2$) | Solid Volume($\mu m^3$) | Total Volume($\mu m^3$) | $S_{inner}/V_{total}$ ($\mu m^{-1}$) | $S_{inner}/S_{outer}$ (/) |
|---|---|---|---|---|---|---|---|
| Max | Health | 1.5158 | 7.2926 | 0.1404 | 0.0429 | 31.9031 | 4.8111 |
| | Damage | 2.8088 | 22.3473 | 0.3198 | 0.0839 | 44.8386 | 9.4213 |
| Min | Health | 0.1903 | 0.2924 | 0.0064 | 0.0038 | 22.7027 | 1.3285 |
| | Damage | 0.1655 | 0.2678 | 0.0064 | 0.0039 | 28.0507 | 1.1687 |
| Median | Health | 0.6911 | 1.7009 | 0.0322 | 0.0254 | 26.1778 | 2.6399 |
| | Damage | 0.7687 | 3.4310 | 0.0839 | 0.0603 | 35.7444 | 4.9164 |
| Mean | Health | 0.6911 | 2.0784 | 0.0428 | 0.0332 | 26.9676 | 2.6784 |
| | Damage | 0.7689 | 5.3469 | 0.0838 | 0.0603 | 35.7444 | 4.6921 |
| SD | Health | 0.3894 | 1.7496 | 0.0344 | 0.0272 | 2.4322 | 0.8903 |
| | Damage | 0.6065 | 4.9664 | 0.0684 | 0.0471 | 4.6908 | 1.6177 |

**Supplementary Table 2** Patient#2 dataset mitochondria properties.

| | | Outer Membrane Surface($\mu m^2$) | Inner Cristae Surface($\mu m^2$) | Solid Volume($\mu m^3$) | Total Volume($\mu m^3$) | $S_{inner}/V_{total}$ ($\mu m^{-1}$) | $S_{inner}/S_{outer}$ (/) |
|---|---|---|---|---|---|---|---|
| Max | Health | 2.1812 | 10.1405 | 0.1562 | 0.2213 | 50.1888 | 5.3391 |
| | Damage | 1.2738 | 4.6058 | 0.0887 | 0.1383 | 38.1473 | 5.2326 |
| Min | Health | 0.1645 | 0.2523 | 0.0070 | 0.0090 | 28.0509 | 1.5342 |
| | Damage | 0.1840 | 0.1998 | 0.0065 | 0.0093 | 21.5140 | 0.8236 |
| Median | Health | 0.8409 | 2.8681 | 0.0508 | 0.0742 | 37.8276 | 3.2750 |
| | Damage | 0.5028 | 1.4345 | 0.0307 | 0.0444 | 30.3534 | 2.4778 |
| Mean | Health | 0.9074 | 3.2610 | 0.0551 | 0.0805 | 38.5514 | 3.3092 |
| | Damage | 0.6338 | 1.7720 | 0.0367 | 0.0549 | 30.5196 | 2.5047 |
| SD | Health | 0.4831 | 2.3745 | 0.0355 | 0.0522 | 4.6909 | 1.0312 |
| | Damage | 0.3332 | 1.3236 | 0.0248 | 0.0380 | 4.0617 | 1.0714 |

**Supplementary Table 3** Patient#3 dataset mitochondria properties.

| | | Outer Membrane Surface($\mu$m$^2$) | Inner Cristae Surface($\mu$m$^2$) | Solid Volume($\mu$m$^3$) | Total Volume($\mu$m$^3$) | $S_{inner}/V_{total}$ ($\mu$m$^{-1}$) | $S_{inner}/S_{outer}$ (/) |
|---|---|---|---|---|---|---|---|
| Max | Health | 1.9337 | 9.1364 | 0.1424 | 0.2731 | 46.2157 | 4.9189 |
| | Damage | 1.0163 | 3.6909 | 0.0608 | 0.0984 | 37.5123 | 3.6316 |
| Min | Health | 0.1756 | 0.2856 | 0.0049 | 0.0077 | 25.7038 | 1.3429 |
| | Damage | 0.1230 | 0.1691 | 0.0044 | 0.0072 | 20.6485 | 1.0220 |
| Median | Health | 0.7545 | 1.9638 | 0.0391 | 0.0633 | 31.6204 | 2.5707 |
| | Damage | 0.3914 | 0.5913 | 0.0131 | 0.0215 | 27.0394 | 1.5458 |
| Mean | Health | 0.7989 | 2.3939 | 0.0485 | 0.0721 | 32.8431 | 2.7349 |
| | Damage | 0.4207 | 0.8396 | 0.0183 | 0.0281 | 27.6619 | 1.7552 |
| SD | Health | 0.4184 | 1.8659 | 0.0324 | 0.0522 | 5.1164 | 0.8543 |
| | Damage | 0.2280 | 0.8147 | 0.0134 | 0.0219 | 4.8590 | 0.6186 |
